# Metabolic Repression of Autophagy Drives Inflammation, Pain, and Malodor in Hidradenitis Suppurativa

**DOI:** 10.64898/2026.01.26.700194

**Authors:** Mahendra P. Kashyap, Rajesh Sinha, Safiya Haque, Lin Jin, Tiffiny Mayo, Craig A. Elmets, Chander Raman, Mohammad Athar

## Abstract

Hidradenitis suppurativa (HS) is a debilitating and underdiagnosed inflammatory skin disease with limited therapeutic options due to an incomplete understanding of its molecular and immunopathogenic basis. Using an integrated multi-omics approach, we delineate convergent pathways linking dysregulated autophagy, malodour, and pain in HS. Lesional tissues exhibited sustained autophagy repression, marked by increased expression and activation of mTOR and ZKSCAN3, reduced levels of autolysosome-promoting metabolites β-hydroxybutyrate and nicotinamide riboside, and accumulation of fructose-1,6-bisphosphate, a negative regulator of AMPK. Transcriptomic analyses identified cadaverine derivatives as key metabolic drivers of keratinocyte reprogramming, inducing gene networks associated with hyperproliferation, inflammation, fibrosis, nociception, and the characteristic carrion-like odour of HS. Concordantly, ATAC-seq revealed increased chromatin accessibility at loci encoding TRP channels and histamine receptors, consistent with heightened nociceptive signalling. Multi-layered proteomics, phospho-proteomics, kinomics, and spatial proteomics analyses, validated by high-resolution confocal imaging, demonstrated elevated mTOR (Ser2448), ZKSCAN3, and P62, alongside reduced LAMP1, across keratinocytes and immune cell populations including CD4⁺ T cells, CD56⁺ NK cells, and CD68⁺ macrophages. Inhibition of mTOR normalized transcriptional programs and cytokine production linked to NLRP3 inflammasome, restoring tissue homeostasis. Collectively, these findings identify a metabolically reinforced mTOR–ZKSCAN3 axis in autophagy dysregulation and cadaverine-driven epithelial reprogramming as central mechanisms sustaining inflammation, pain, and fibrosis in HS.

**Highlights:**

- We identified BCAAs, BHB, NR, and FBP as key metabolites linked to mTOR activation and autophagy dysregulation in HS.
- Polyamines like cadaverine and putrescine, their acetylated derivatives, LPCs, and endocannabinoid dysregulation likely drive malodor and pain in HS.
- Cadaverine treatment of Ker-CT cells induces HS-associated gene signatures of inflammation, fibrosis, and pain.
- Identification of an HS macrophage subpopulation expressing ZKSCAN3 with activated mTOR and impaired autophagy.

## INTRODUCTION

Hidradenitis Suppurativa (HS) is a chronic, debilitating inflammatory skin disease of poorly defined pathogenesis ^1,2^. HS patients experience restricted physical function, social isolation, diminished self-esteem, and impaired intimacy ^3^. HS comorbidities include metabolic syndrome, obesity, type-2 diabetes, hypertension, myocardial infarction and depression often with increased risk of suicide ^4^.

Despite the availability of FDA-approved biologics, including anti-TNFα (adalimumab), anti-IL-17A (secukinumab), and dual IL-17A/F inhibitor (bimekizumab), clinical responses in HS remain unsatisfactory ^1,5^. This limited efficacy underscores the involvement of additional undefined pathogenic mechanisms and highlights the urgent need for a deeper mechanistic understanding of HS to guide the development of novel therapies. In addition, lack of an animal model that faithfully recapitulates the clinical and immunological features of HS, restricts mechanistic studies and therapeutic discovery.

Previously, we demonstrated that aberrant protein translation signaling drives hyperproliferation of follicular epithelial cells in HS, a pathological feature associated with an elevated risk of squamous neoplasm development ^6^. Consistently, Adawi *et al*. showed the transcriptomic profile of tunnel epithelium more closely resembling squamous cell carcinoma than infundibular cysts ^7^. More recently, we identified a heterogeneous population of epithelial progenitor cells that overexpress S100A7/8/9 and KRT6 family members, which actively regulate IL-1 and IL-10–mediated signaling as well as complement cascade–driving inflammatory responses in HS ^8^. We further uncovered that CD2–CD58 cognate interactions among innate and adaptive lymphoid cells, epidermal cells, and fibroblasts play a critical role in driving the inflammatory pathogenesis of HS ^5^. Here, we further focused on metabolic basis of this disease due to its known association with metabolic comorbidities including obesity, diabetes etc. In this context, dysregulated mTOR signaling and impaired autophagy–key regulators of cellular homeostasis and hallmarks of inflammatory and metabolic disorders ^9^ –may represent a central mechanism driving hyperinflammation in HS. Loss or disruption of autophagy-related genes impairs immune cell differentiation, fuels excessive ROS production, and drives inflammasome activation, collectively amplifying tissue damage and inflammation ^9^. Recently, inflammasome activation has emerged as a player in the pathogenesis of HS ^10^. Importantly, GLP-1 agonists, which are potent anti-diabetic and anti-obesity drugs, are known to stimulate autophagy ^11^ and significantly reduce severity of HS in a recent case study ^12^.

Here, we integrate comprehensive metabolic landscape profiling with multi-omics approaches to elucidate the molecular circuitry driving HS pathogenesis. We uncover key metabolites and signaling nodes that converge on mTOR-dependent autophagy dysregulation across hyperproliferative keratinocytes, ZKSCAN3⁺ macrophages, and CD56⁺ NK cells, revealing how these cell populations cooperatively shape the inflammatory and tissue-remodeling landscape of HS. Furthermore, reduced levels of the autolysosome-promoting metabolites β-hydroxybutyrate (BHB), nicotinamide riboside (NR) and elevated fructose-1,6-bisphosphate (FBP), a metabolic inhibitor of AMPK signaling play critical role in sustaining metabolic alterations both in immune and non-immune cell populations. Transcriptomic analyses revealed that treatment of keratinocytes with cadaverine, a malodor-associated polyamine, drives keratinocyte gene programs that promote hyperproliferation, pro-inflammatory and fibrotic signaling, and nociceptive responses. While ATAC–seq profiling of HS keratinocytes revealed increased chromatin accessibility at loci encoding nociceptive transient receptor potential (TRP) channels and histamine receptors in HS. These data provide a molecular basis for metabolic regulation leading to key histopathological and phenotypic characteristics of HS.

## RESULTS

### Metabolic landscape uncovers key drivers of oxidative stress, malodor, pain, and autophagy dysfunction in HS

Comprehensive metabolomic profiling of HS (n=14) and healthy control (n=10) skin biopsies using UPLC-MS/MS on the global HD4 dataset, identified 657 biochemicals. Of these, 615 compounds were known (named biochemicals), and 42 compounds were with unknown structural identities (unnamed biochemicals). Briefly, 311 biochemicals were increased and 42 were decreased, significantly in HS skin relative to normal skin (Figure 1A). The PCA plot of the global HD4 dataset shows clear separation along the first principal component, highlighting a significant metabolomic distinction between HS and normal skin samples (NS) (Figure 1B). The lipidomic platform identified a total of 959 lipid metabolites of known identity (named biochemicals) in the complex lipid profile (CLP) dataset. Of these, 215 lipid metabolites were upregulated, while 5 were downregulated, significantly in HS (Figure 1C). The PCA plot of the CLP dataset reveals clear separation along the second principal component (Figure 1D). Heatmaps of metabolomics (HD4 dataset) reveal significant alterations in metabolite profile between HS and NS (Figure 1E). Most altered metabolites map to pathways governing antioxidant signaling, energy production, amino acid and nucleotide metabolism, protein turnover, autophagy, lipid metabolism and pain perception (Figures 1, 2 and S1).

**Fig 1.**
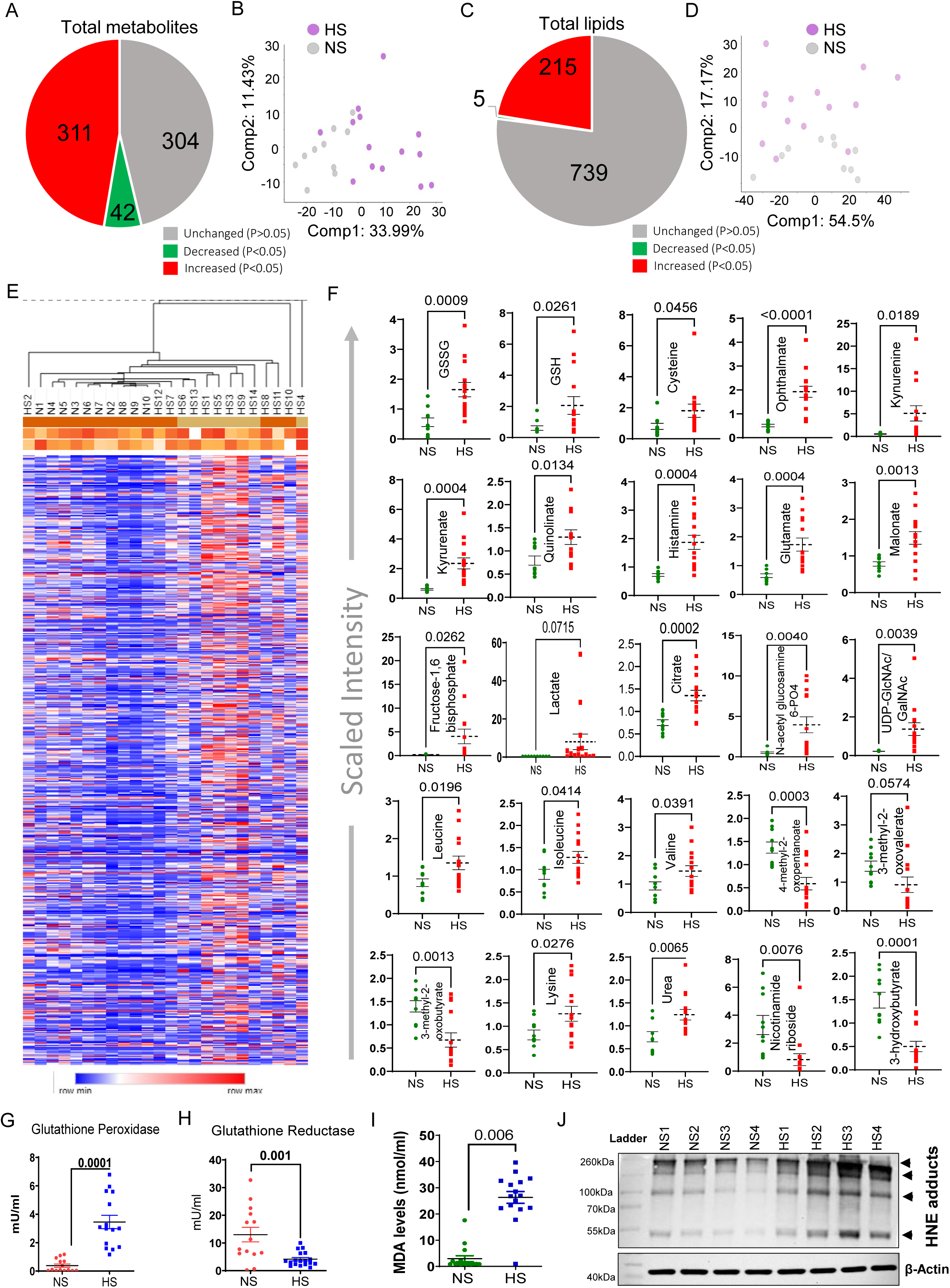
Cutaneous Metabolic profiling of HS and normal skin. HS (purple dots) vs NS (grey dots) analysis of biochemicals: **(A)** Pie chart **(B)** PC on HD platform (C) pie chart and (D) PC on CLP platform. **(E)** Heatmap shows expression levels of named metabolites in HS (n=14) and normal skin (n=10). The color scale indicates relative level of the abundance of each metabolite with red indicating levels greater than the mean value and blue indicating levels below the mean value. **(F)** Graphs showing significantly altered metabolites involved in various metabolic pathways in HS skin vs normal skin. **(G-H)** Graphs showing the altered activity of glutathione peroxidase and glutathione reductase in HS. **(I)** MDA (malondialdehyde) levels in HS and normal skin. Dots in each graph represent individual HS vs NS. **(J)** Western blot analysis of HNE adducts in HS and NS.

In HS skin, glutathione metabolism is significantly altered, as reflected by elevated levels of reduced glutathione (GSH), oxidized glutathione (GSSG), the rate-limiting amino acid cysteine, and ophthalmate — a derivative of glutathione (Figure 1F). Notably, the higher oxidized-to-reduced glutathione ratio in HS skin (1.43±0.24) relative to NS (0.92±0.19) underscores a pronounced oxidative environment (Figure S2A). The reduced activity of glutathione reductase along enhanced activity of glutathione peroxidase further validates these metabolomics findings (Figures 1G and 1H). Elevated levels of malondialdehyde (MDA) (Fig. 1I) and HNE-protein adducts along with enhanced expression of oxidative stress-responsive proteins; NRF2 and HO-1 further confirmed strong oxidative environment in HS (Figures 1J and S2B).

High levels of quinolinate (QA or Quin, 2,3-pyridine dicarboxylic acid; 241.89-fold; p<0.00) along with elevated histamine, kynurenine and kynurenate in HS skin, indicate hyperinflammatory immune environment in HS skin (Figure 1F). Guenin-Mace *et al.*^13^ also reported upregulated levels of quin, kynurenine and expansion of quin^+^CD45^+^ cells in lesional HS skin. Histamine is a vasodilator while kynurenine is generated from tryptophan via indoleamine 2,3-dioxygenase, which can be activated by IFN-γ and TNF-α^14^ cytokines, which are elevated in HS. Dysregulated tryptophan catabolism is also found in the host skin-microbiota interface in HS ^13^. In neuronal tissues, QA works as a N-methyl-d-aspartate (NMDA) agonist by potentiating Mg^2+^-sensitive NMDA receptors which allows enhanced Ca^+2^ flux to enter the cell leading to the activation of various endonucleases, phospholipases, and proteases ^15^. These molecular events are critical to compromise cellular integrity. Moreover, QA also increases the cell’s response to pain stimuli ^16^. Glutamate, an activator of NMDA receptors, is significantly elevated in HS skin (Figure 1F). Thus, sensitization of nerve terminals by these chemicals, may lead to debilitating pain as observed in HS ^17^. Activated macrophage are known source of QA production and express high levels of NMDA receptors in the inflamed tissue environment ^18^. We also observed a significant increase expression of inflammatory survival factor, GRINA in CD68^+^ macrophages^5^ (Figure S3), reinforcing the enrichment of proinflammatory macrophages in HS lesions ^19^. Earlier, we reported a striking abundance of CD38^+^ inflammatory macrophages in lesional HS skin ^20^ which is an enzyme that act via consumption of NAD for immune cell activation, proliferation and function. The levels of NAD+ are elevated in HS skin by 1.63-fold (Figures S1C).

Activation of innate immune cells relies heavily on glycolysis, while the TCA cycle is typically associated with the inflammation-resolving phenotype ^21^. The significantly elevated levels of 3-phosphoglycerate, fructose-1,6-bisphosphate (FBP), phosphoenolpyruvate, and lactate, the products of glycolysis confirm enhanced glycolytic activity in HS skin (Figures 1F and S1D). FBP levels were inversely correlated with AMPK activation which acts as an autophagy inducer ^22^. Therefore, high levels of FBP accumulated in HS lesions correlate well with AMPK inhibition (Figure 4D and discussed in subsequent sections of this manuscript). Autophagy inhibition facilitates the secretion of IFNs type I, II and TNF-α *in vitro* and *in vivo* ^23,24^. Recently, treatment with metformin, an inducer of AMPK, suppressed the expression of key glycolytic genes and reduced the production of various cytokines/chemokines in peripheral blood mononuclear cells from HS patients ^25^.

Branched-chain amino acids (BCAAs), including leucine, isoleucine, and valine, are key substrates for energy production to fuel the TCA cycle. In HS, we observed elevated levels of these three essential BCAAs, alongside decreased levels of their respective catabolites, such as 4-methyl-2-oxopentanoate, 3-methyl-2-oxovalerate, and 3-methyl-2-oxobutyrate (Figure 1F). These alterations suggest increased protein breakdown in HS, resembling the metabolic state seen during fasting ^26^. BCAA catabolism is vital for energy generation, particularly in obese individuals, and elevated BCAA levels are associated with the development of type 2 diabetes ^27,28^. Indeed, these are known co-morbidities associated with HS ^29^. About 75% of BCAAs are recycled into protein synthesis, remaining are oxidized for energy generation. This is consistent with elevated dipeptides in HS, indicating enhanced protein turnover (Figures S1E and S1F).

BCAAs also activate mTORC1 ^30^. We showed enhanced levels of eIF4F complex in HS, which regulates 5′-cap-dependent protein translation ^6^. In this regard, mTOR facilitates phosphorylation-dependent activation of translation elongation factor eIF4E, which we have shown to be overexpressed in HS skin ^6^. Elevated levels of nucleotide sugars such as UDP-GlcNAc and UDP-GalNAc and with their precursors, glucosamine 6-phosphate and N-acetylglucosamine 6-phosphate are also observed in HS (Figures 1F and S1D). These sugars serve as donor molecules in glycosylation reactions, during post-translational modifications of proteins which are essential for proliferating cells along with high protein supply ^31^. This aligns with our previous observation of increased cell cycle gene expression in HS skin ^8^.

Increased amino acid catabolism or impaired urea clearance may lead to elevated urea levels as observed in HS skin (Figure 1F). Interestingly, Beclin-1-mediated activation of autophagy has been shown to improve urea cycle disorders ^32^. However, as we discussed later, Beclin-1 levels are significantly reduced in HS skin providing mechanistic support to these data. Another intriguing metabolite, β-hydroxybutyrate (BHB), synthesized during the metabolism of ketogenic amino acids and fatty acids, is decreased in HS skin (Figure 1F). BHB has been shown to mitigate sepsis-associated acute lung injury by promoting autophagy in macrophages ^33^ with known role in attenuating NLRP3 inflammasome ^34^. BHB also inhibits class I histone deacetylases (HDACs), regulating the expression of critical genes such as NF-κB, TP53, MYC, and MYOD ^35^. This aligns with our earlier findings showing high expression of c-Myc in HS-associated keratoacanthoma ^6^.

Here in HS lesions, we report the augmented synthesis of malodorous chemicals such as diamine cadaverine (15.53-fold; P≤.0208), putrescine and their acetyl derivatives N-acetyl cadaverine and N-acetyl putrescine along with elevated polyamine spermine (Figures 2A and 2B), which uncovers the undescribed important phenotype of this disease. N (‘1)-acetyl spermidine, which is elevated in HS, is also known to be oxidized into putrescine. Polyamines facilitate cell growth and development by regulating translation initiation factor 5A (eIF5A) ^36^. Earlier, we have shown that elevated ornithine decarboxylase that augment polyamine production plays a significant role in the pathogenesis of skin carcinogenesis ^37^. Elevated levels of polyamines in HS skin are also consistent with the known enhanced risk of these lesions for developing squamous neoplasm ^6^.

**Fig 2:**
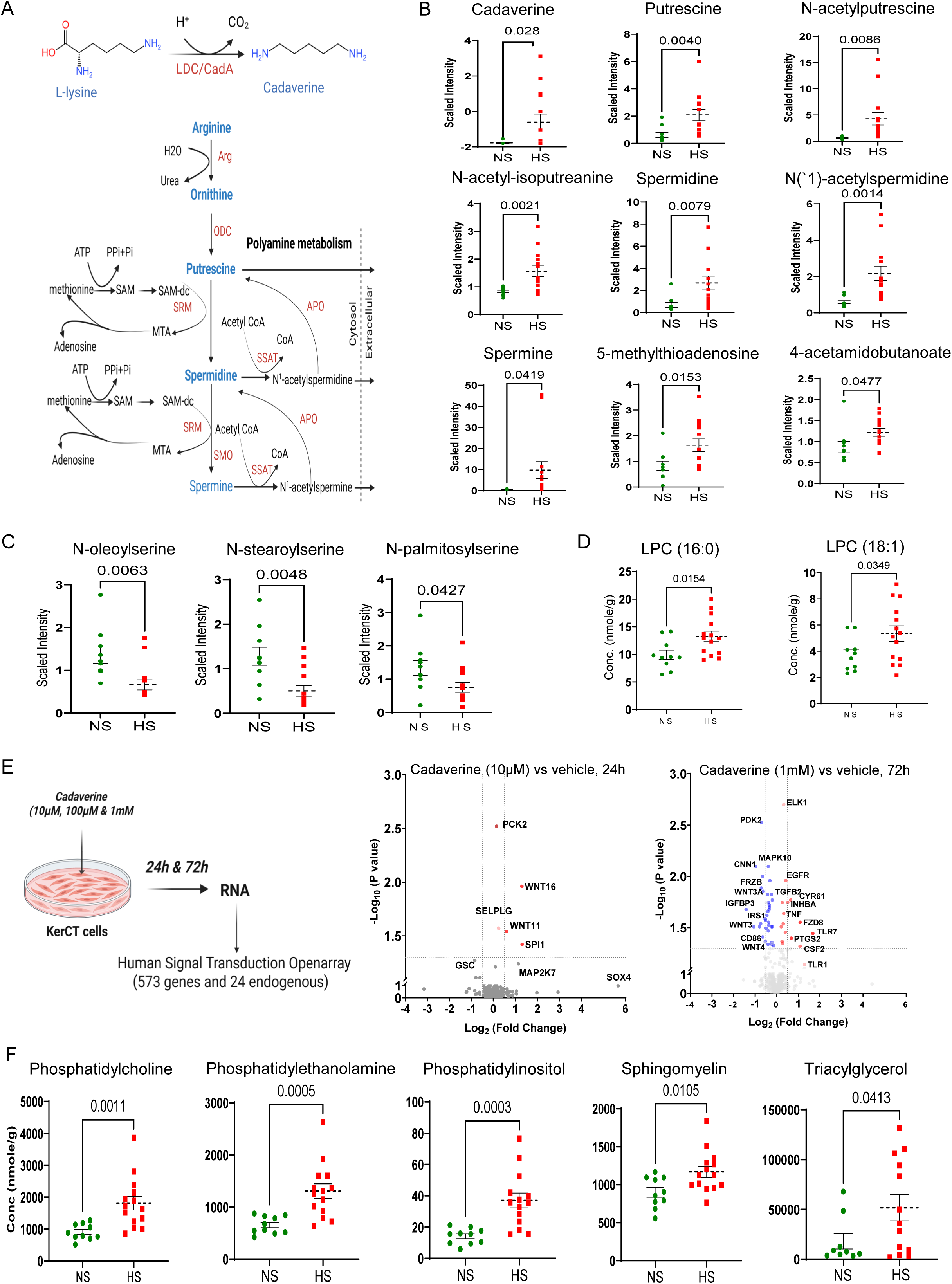
Altered metabolites involved in Malodour, pain and lipid signalling. **(A)** Schematic diagram illustrates polyamine metabolism. **(B)** Graphs showing the significantly altered diamines, polyamines and their derivates involved in malodours pathways from HS. Graphs showing significantly altered **(C)** endocannabinoids and **(D)** lysophosphatidylcholines in HS. **(E)** Diagram (left) shows experimental approach of cadaverine exposure on Ker-CT cells and volcano plots showing genes involved in signal transduction pathways in Ker-CT cells in response to cadaverine challenge at 10 µM for 24h and 1mM for 72h. **(F)** Graphs showing levels of phosphatidylcholine (PC), phosphatidylethanolamine (PE), phosphatidylinositol (PI), sphingomyelin (SM) and cholesteryl ester (CE) in HS vs normal skin. Dots in each graph represent individual HS vs NS.

Polyamines are known to be associated with TRPV1-mediated nociception in inflammatory diseases ^38^ and their elevation in HS along with down-modulated endocannabinoids, which are known to suppress the perception of neuropathic pain via activation of cannabinoid receptor CB1 ^39^, may serve as an underpinning mechanism of sustained pain in HS patients. HS skin exhibited a profound shift in lipid metabolism, characterized by reduced endocannabinoids (N-oleoyl serine, N-stearoyl serine, and N-palmitoylserine) (Figure 2C) alongside elevated pro-inflammatory lysophosphatidylcholine species (LPC 16:0 and LPC 18:1) (Figure 2D). Emerging evidence suggests that lysophosphatidylcholines function as direct mediators of neuropathic pain ^40,41^.

Cadaverine and putrescine, which typically arise from bacterial decarboxylation of the basic amino acids, ornithine and lysine, are known for strong and repulsive odor to human ^42^. Thus, the carrion odor in HS patients may be associated with production of these metabolites in lesional skin associated with bacterial or microbial infection. To directly assess the importance of malodorous metabolites in HS, we treated normal human keratinocytes (Ker-CT) with cadaverine under controlled *in vitro* conditions (Figures 2E and S4-S5). Interestingly, even low dose of cadaverine (10µM) at early time point (24h) induced the expression of transcription factor *SPI1* (PU.1) (Figure 2E), which has known role in psoriasis where it modulates the expression of inflammatory genes (*IL23 and TNFA*) in macrophage ^43^. Additionally, cadaverine at a higher dose (1mM) activates the transcription factor SOX4 (Figure S4C), which regulates epithelial–mesenchymal transition, CXCL13 production, and squamous cell carcinoma ^44–46^. Strikingly, treatment with the highest dose of cadaverine (1mM) for extended period (72h) leads to upregulation of several inflammatory drivers: *TNFA, TLR7, CSF2*; pain sensor *PTGS2 (COX2)*; fibrotic and angiogenic markers such as *TGFB2, CYR61 and EGFR*. Cadaverine-induced elevated levels CSF2/GM-CSF and downregulations of *IRS1* in this study may contribute to the fibrotic skin lesions in HS (Figure 2E). Moreover, unbiased IPA data revealed inhibition of AMPK signaling [Z score= -2.236; -log(p-value) = 6.33)], while activation of mTOR signaling [Z score= 0.95; -log(p-value) = 1.10)] by cadaverine which mimics accumulation of this metabolite in chronic HS lesions (Figure S5D).

While several metabolites belonging to purine and pyrimidine metabolism, pregnenolone and androgenic steroids metabolism, and primary bile acid metabolism were significantly upregulated, it is particularly noteworthy that key metabolites involved in nicotinate and nicotinamide metabolism were significantly downregulated In HS (Figure S6). Nicotinamide riboside (NR; 0.47-fold; p≤0.0001) and nicotinamide ribonucleotide (NMN; 0.47-fold; p≤0.0017), which promote autolysosome clearance, were downregulated in HS skin (Figure 1F and S5C). Boosting NAD+ levels by NR ameliorates TLR4-mediated type I IFN in systemic lupus erythematosus monocytes ^47^. Interestingly, nicotinamide and zinc supplementation show some benefit as maintenance therapy in mild to moderate HS patients who have previously been treated with minocycline ^48^.

Thus, reduced levels of BHB, NR and elevated levels of FBP as well as abundance of various amino acids in HS corroborate autophagy inhibition (Figure S7).

Many phospholipids such as phosphatidylcholine, lysophosphatidylcholine, sphingomyelin, and ceramide, which serve as building blocks for cells and autophagy for autophagy vacuole membrane formation, were elevated in HS. However, enhanced synthesis of fatty acids as reflected by elevated malonate, triglycerides and cholesterol ester (Figures 1F, 2F and S8-S9), point to autophagy dysregulation ^49^. Autophagy dysregulation is also apparent from the IPA of metabolomics data showing downregulated sirtuin signalling pathways, enhanced amino acid levels and downregulation of LDHA (-log(p-value) = 2.21E-08, z-score= -2.828). CD40, which is mainly expressed on antigen-presenting cells (APCs) and B cells, was rank 1 upregulated (-log(p-value) = 1.06E-12, Z-score= 3.771) (Figure S10).

### Global proteomic profiling highlights mechanisms driving inflammation, pain, and autophagy dysregulation in HS

To strengthen the molecular understanding of HS pathogenesis, we carried out proteomics/phospho-proteomics profiling of HS (n=5) and NS (n=5). Total, 15081 peptides were visualized, of these 13326 peptides passed quality control. 3535 proteins identified through the database search output filtered to 1% FDR (False Discovery Rate). 980 proteins were differentially expressed in HS tissue of which 307 proteins were upregulated (FC > |1.5|, p-value<0.05) while 54 proteins were down-modulated (FC < |-1.5|, p-value<0.05).

Data showed downregulation of several collagens with concurrent upregulation of collagenases, suggesting excessive extracellular matrix (ECM) degradation. Elevated levels of multiple small proline-rich (SPRR) family proteins in HS skin indicate enhanced cornified envelope formation in keratinocytes, reflecting aberrant epidermal differentiation ^50^ and compromised barrier integrity, which may promote follicular occlusion and reinforce inflammatory and fibrotic remodeling. Additionally, increased levels of MAPKs, calpain, NFκB1, and STAT family proteins were observed (Figures 3A and 3B). Enhanced collagen degradation is a hallmark in various inflammatory diseases, including rheumatoid arthritis, epidermolysis bullosa, and periodontal diseases ^51^. Macrophage related proteins such as macrophage-capping protein (CAPG), macrophage mannose receptor 1 (MRC1), macrophage colony-stimulating factor 1 receptor (CSFR1), and macrophage migration inhibitory factor (MIF) were elevated in HS skin (Figure 3A). MIF negatively regulates P53 activity, thus affording protection to macrophages cell death in a hyper-inflammatory environment ^52^ which is consistence with our observation of increased expression of GRINA. Another crucial finding in proteomics data was upregulation of ZKSCAN3 along with downmodulation of several fundamental subunits of mitochondrial ATP synthetase (Figure 3A). Confocal imaging established the cellular compartmentalization of ZKSCAN3 prominently in keratinocytes and prominently in CD68^+^ macrophages (Figures 3C and S11). ZKSCAN3 is considered as a master transcriptional repressor of autophagy genes ^53^ while ATP synthetase is required for energy generation for autophagy ^54^. For example, *P. aeruginosa* infected *Zkscan3*^−/−^ mice manifest enhanced autophagy-lysosomal pathway (ALP) and bacterial clearance ^55^. Consistently, *ZKSCAN3* overexpression reduces rapamycin-induced autophagy in various cell types ^53^. Therefore, we consider that ZKSCAN3 as a critically important protein in tempering autophagy in HS.

**Fig 3.**
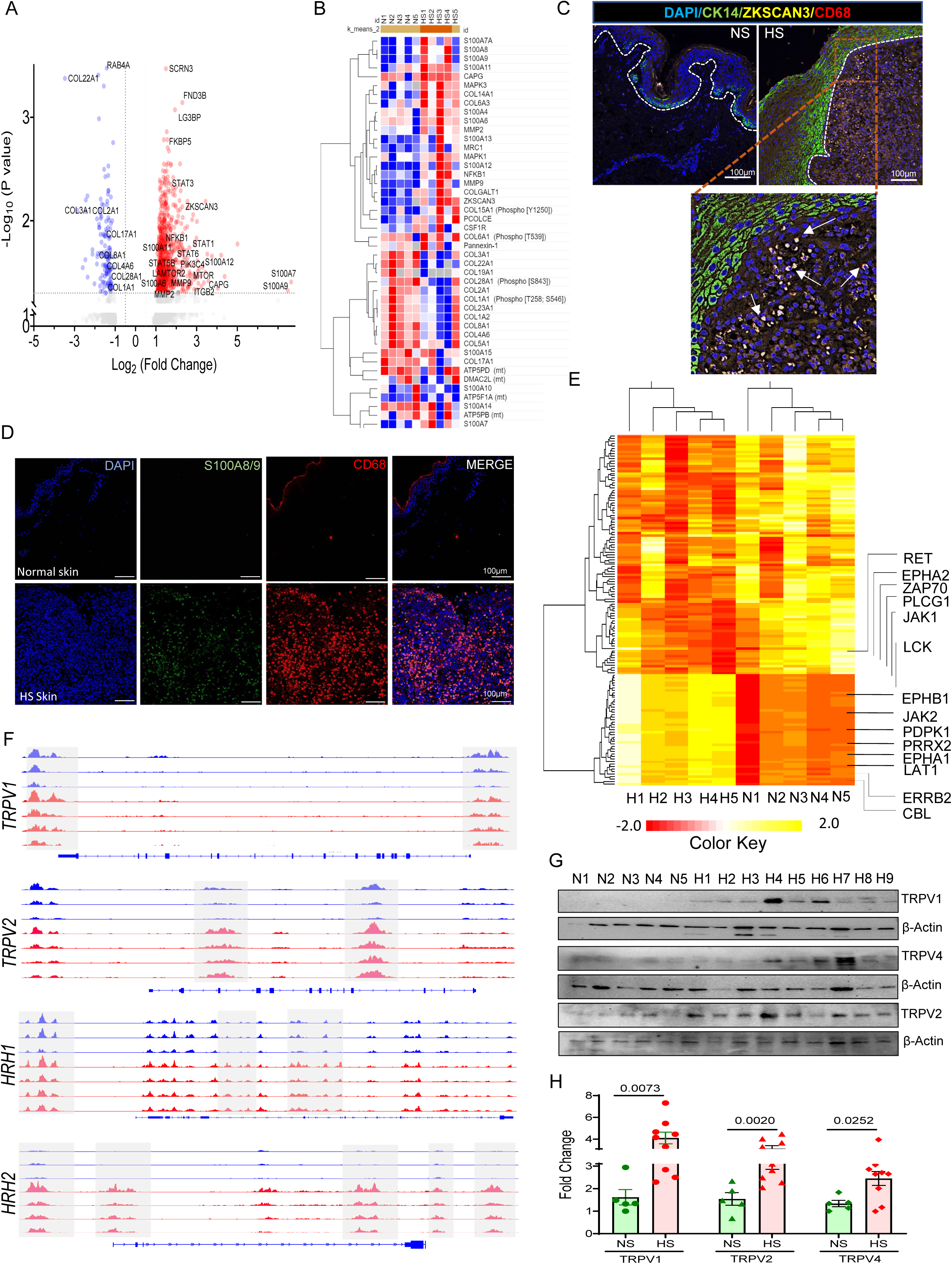
Proteomics and kinomics profiling of HS and normal skin. **(A)** Volcano plots of proteomics showing differentially expressed proteins in HS vs NS. **(B)** Heatmap shows expression difference in proteins of S100 family, collagens and collagenase, ZKSAN3, and subunits of mitochondrial ATP synthetases. **(C)** Representative confocal micrographs showing expression of cytokeratin-14 (CK14, green), ZKSCAN3 (yellow) and CD68 (Red) IN NS and HS. **(D)** Confocal micrographs showing enhanced expression of S100A8/A9 heterodimers (green) in CD68^+^ (red) macrophages in HS vs NS. **(E)** Heatmap showing differences in kinase activity profile between HS (n=5) and NS (n=5). Yellow denotes high activity while red reflects lower enzymatic activity. **(F)** The integrative Genomics Viewer (IGV) Genome browser representative view of indicated gene loci involved in pain - *TRPV1*, *TRPV2*, *HRH1* and *HRH2* characterized by enrichment in H3K4me1 (dot boxes) in healthy vs. HS CD49fhigh basal cells (n=3 for healthy, n-4 for HS). **(G)** Western blots and **(H)** densitometry analysis of TRPV1, TRPV2 and TRPV4 receptors in HS (n=9) vs NS (n=5).

Phospho-proteomics data identified that phosphorylation of 27 proteins was significantly altered in HS. Out of 27, 19 proteins had significantly enhanced phosphorylation, however, 8 proteins had significant reduction in the phosphorylation. Heatmaps of proteomics data showed high abundance of alarmins namely, S100A6, S100A7/S100A7A, S100A8, S100A9, S100A12 and their downstream targets ERK1/2 and NFκB1 (Figure 3B). Consistently, we also observed high levels of pERK and pNFΚB proteins in HS skin (Figure S12A). Earlier, we reported upregulated S100A8, S100A9, MYD88 along with downstream effector NFΚB1 at single cell resolution level in HS ^5^. Confocal data confirms high expressions of S100A8/A9 heterodimer in CD68^+^ macrophages in HS Skin (Figure 3D). Various proinflammatory mediators like GM-CSF, TNF-α, IL-1β and LPS, which are elevated in HS, induce the release of S100A8/A9 from monocytes ^56^. Recently, we also described the role of S100 family proteins in inducing plasticity in HS epithelial cells ^8^.

Secernin-3 (SCRN3) with a role in thermal nociception ^57^, was overexpressed in HS (Figure 3B). This protein is induced in macrophage in response to LP challenge (Figure S12B).

### Kinomics and ATAC-Seq uncover immunopathogenesis of HS-associated pain

We further complemented our phospho-proteome data by kinomics which measure activity of 196 tyrosine peptide substrates and 144 serine/threonine kinase peptide substrates using PamGene microarray platform. Phospho-peptides associated with kinases involved in autophagy inhibition and neuropathic pain include RET^58^, ERBB2^59^, PDPK1 and Ephrin receptor kinases such as EphA2 ^60^ showed enhanced activity in HS (Figure 3E). Importantly, EphA was found to be associated with the neuropathic pain by activating CXCR4/RhoA/ROCK2 pathway in animals having chronic constriction injury (CCI)^5,61^consistent with our earlier data and a recent publication^62^, this pathway may be involved in the pathogenesis of neuropathic pain in HS. Our ATAC-sequencing (ATAC-seq) data also identified open chromatin pattern at gene bodies of various TRP and histamine receptors namely, *TRPV1, TRPV2*, *TRPV4*, *HRH1* and *HRH2* in HS epithelial cells relative to normal keratinocytes (Figure 3F). Western blot analysis further affirms upregulated protein expression of TRPV1 and TRPV4 in HS skin (Figures 3G and 3H).

Other kinases, such as zeta-chain-associated protein kinase-70 (ZAP70), CBL, LCK, PRRX2, PGFRB, and fibroblast growth factor receptors namely, FGFR2 and FGFR3 that were found to be activated, are known to control the NK cells, T cells, and fibroblast activation. Phosphorylation of Zap-70 in NK cells increase granzyme-B (GZM-B) expression and NK-cell activity triggered by the immunomodulatory drugs (IMiDs)^63^. Earlier, we have demonstrated that IKZF1 inhibitor lenalidomide, an IMiD, reduces cytokine/chemokine secretion in HS organotypic cultures ^64^. LCK, is central to signal transduction downstream of the TCR and NK cell-activating receptors ^65^. We recently uncovered the role of NK, NKT and CD4T cells in HS pathogenesis ^5^. LAT1, an L-type amino acids transporter, was upregulated (Figure 3F) which is consistent with mTOR activation. Our, proteomics and phospho-proteomics data concur with these kinomics data to establish an intricate mechanism of receptor tyrosine kinases, many of these are regulated by PI3K/AKT signaling which activates mTOR, PLCG/PKC, and JAK/STAT pathways and are responsible for secretion of cytokines such as IL6, IL10 or IL22. IPA analysis of kinomics confirmed the induction of IL-15 production, RAF/MAPK cascade, NK cell signaling, PI3K/AKT signaling, Insulin receptor signaling, FGF signaling while inactivation of PD-1, PD-L1 cancer and PTEN signalling (Figures S13A and S13B). Overall, these data provide a cross validation and rigor of our multi-omics approaches and confirm a link for dysregulated autophagy and metabolism in the pathogenesis of malodourous and highly painful lesions in HS.

### High-Resolution Spatial Proteomics and Confocal imaging Reveals Cell-Type–Specific Autophagy Dysregulation in HS Lesions

To advance observations of dysregulated autophagy, we further delineate the localization and role of autophagy dysregulation in HS. Unbiased high-resolution CosMx analysis of HS (n=7) and NS (n=3) FFPE slides which enabled 68 proteins at a single-cell resolution revealed that LAMP1 protein expression was reduced in tunnel-associated epithelium compared with the epidermis within the same lesion. Interestingly, LAMP1 expression was further reduced in CD4⁺ (yellow), CD56⁺ (red) and CD68⁺ (green) cells (Figures 4A and S14), indicating impaired autophagy in these immune populations. This observation is consistent with our previous scRNA-seq analysis, which revealed downregulated mRNA expression of the autophagy-related protein MAP1LC3B in the similar immune cell populations ^5^ (Figure S15). These data suggest compromised lysosomal functions in tunnel-associated epithelial, and majority of tissue localized immune populations. ALP is required to ensure sustained autophagy flux and autophagosome lysosome fusion that generates autolysosome to degrade cellular debris. Epidermis adjacent to tunnel region showed higher LAMP1 expression in epithelium (Figure 4A). Consequently, levels of IL18 (blue) not IL1β (magenta) were high in tunnel-associated epithelium which showed robust immune infiltration in the tunnel region (Figures 4B and S15). Our previous scRNA-seq analyses revealed that IL18 expression is largely restricted to epithelial cell ^66^, highlighting epithelized tunnels as a major driver of inflammatory mediators in HS lesions ^67^.

**Fig 4.**
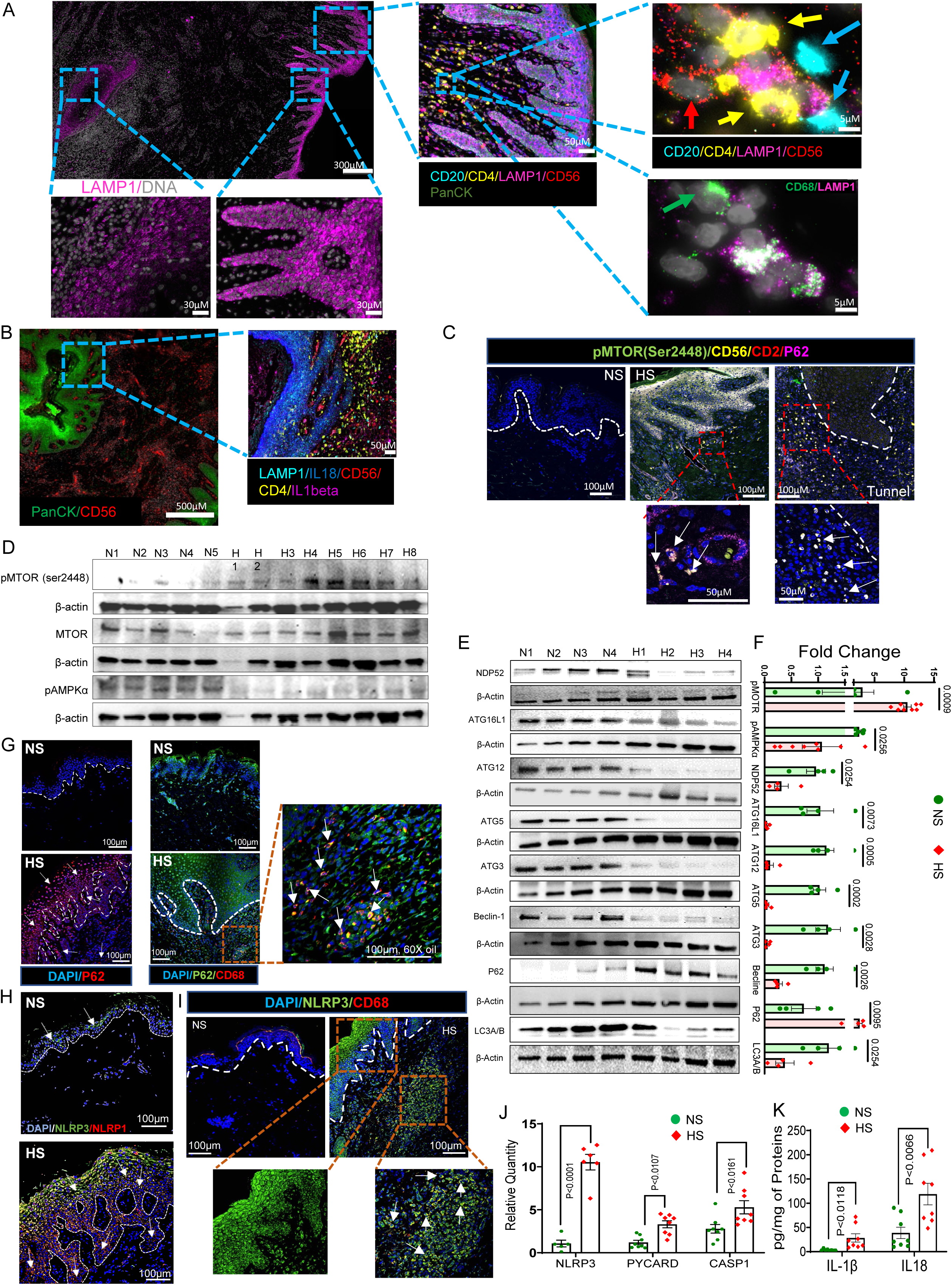
Dysregulated autophagy-dependent NLRP3 inflammasome signaling in HS skin. **(A)** Spatial Proteomic (CosMx) images showing expression of LAMP1 in Tunnel associated epithelium relative to acanthotic epithelium in HS lesions. Elaborated area below acanthotic epithelium shows expansion of CD4 (yellow), CD56 (red), CD20 (aqua) and CD68 (green) cells which lack LAMP1 (magenta). **(B)** CosMx proteome images showing PanCK (green) and CD56 (red) cells in HS skin. Elaborated area of tunnel region showing expression of IL18 (blue) with LAMP1 (aqua) and expansion of CD56 (red) and CD4 (yellow) along with IL1β (magenta). **(C)** Immuno-localization by confocal microscopy of pMTOR (ser2448; green) in the epidermis and CD56+(yellow)/CD2+(red) NK cells in hypodermal area of HS skin. mTOR^+^ NK (CD56+) cells express high levels of P62 (pink). **(D)** Western blots showing enhanced pMTOR (ser2448) while low levels of AMPKα in HS (n=8) relative to normal skin (n=5). **(E)** Western blots (n=4 normal and n=4 HS patients) of various proteins involved in regulation of autophagy. **(F)** Densitometric analysis (bar graph) show fold change in expression of proteins. Data are presented mean ± SEMp; p-values; unpaired Student’s t test. **(G)** Confocal microscopy images demonstrating enhanced accumulation of autophagy receptor P62 (red) throughout the hyperproliferative epithelial compartment. White arrows show expression of P62 in epidermal layers (left panel). P62 (green) expression is elevated in CD68^+^ (red) macrophages in HS Skin (right panel). White arrows in expended inset show expression of P62 in CD68^+^ (red) macrophages **(H)** Immunofluorescence micrographs show NLRP3 (green)/NLRP1 (red) staining across the hyperproliferative epidermis. **(I)** Expression of NLRP3 (green) and CD68 (red) show enhanced activation of NLRP3 inflammasome in CD68^+^ macrophages (white arrow). **(J)** Real time qPCR analysis shows upregulated mRNA expression of *NLRP3, PYCARD* and *CASP1* (n=8 each HS or NS). **(K)** Protein levels of IL1β and IL18 by Luminex assay (n=8/group).

Confocal analysis demonstrates elevated phosphorylation of mTOR (Ser2448) and enhanced staining of P62 in acanthotic epithelial layer and CD56^+^CD2^+^ NK cells in hypodermal region of HS skin (Figures 4C and S16). Western blot analysis validated higher phosphorylation of mTOR (Ser2448) while reduced phosphorylation of AMPKα in HS skin (Figure 4D). As shown in Figures 4E-F, the expression of several autophagy-related proteins namely, NDP52; ATG16L1; ATG12; ATG5; ATG3; Becline-1 and LC3A/B was decreased in HS skin relative to NS. However, autophagy receptor P62, which usually consumed during normal autophagic flux, was accumulated in insoluble fraction of HS tissues. We observed an enhanced staining of P62 throughout the epithelial cells of psoriasiform epidermis (Figure 4G) in HS and CD68^+^ macrophages (Figure 4G). We propose that ZKSCAN^hi^ macrophages with P62 sequestered in insoluble fractions are a critical pathogenic cell population in the immuno-pathogenesis of HS.

Enhanced staining of NLRP3 inflammasome components was seen throughout the epithelium (Figure 4H) and CD68^+^ macrophages (Figure 4I). Dysregulated autophagy is known to activate of NLRP3 ^68^. Upregulated mRNA transcripts of *NLRP3* were also aligned to its downstream target genes *PYCARD*, *CASP1* and *NLRP3* and effector proteins, IL-1β and IL18 in HS (Figure 4J and 4K). Additionally, reanalysis of our scRNA data showed that macrophages in HS lesion express high levels of mRNA of *NLRP3* ^5^. Our CosMx data containing mRNA and protein panels are generated as part of another ongoing study, while we only extracted relevant protein data complementing this multi-omics study.

### Attenuation of mTOR and dampening of inflammasome activity reprogram signal transduction networks driving HS pathogenesis

Our multi-omics data point that mTOR-regulated metabolic switch aberrantly dysregulates protein translation control, autophagy and metabolism leading to malodorous, highly inflamed and painful lesions in HS ^6^. We therefore employed pharmacological modulators targeting mTOR, inflammasomes, and autophagy using rapamycin, MCC950, and 3-methyladenine (3-MA), respectively (Figure 5A). To assess their effects, we employed TaqMan-based inflammatory and signal transduction openarray panels (containing 1183 target genes) using our revalidated skin explant culture model (Description provided in supplemental section and Figure S17). HS skin explants-treated with rapamycin (n=3) or MCC950 (n=3), showed a reversal in expression pattern of inflammatory and signal transduction genes showing a trend towards the normalization of homeostasis (Figure 5B). Volcano plots affirmed that rapamycin significantly downregulated mRNA levels of *MTOR*. Genes involved in pattern recognition receptors (PRRs) signalling such as *NLRP3*, *PYCARD*, alarmins such as *S100A9, S100A12*; TFs such as *JUN, IRF1, GATA3, STAT3, STAT5B*; various cytokines/chemokines and their receptors such as *IL33, IL1R1, IL1R2, IL1RL2, IL1RAP, IL2RG, IL1F7, IL10RA, IL12RB, IL15RA*, *IL18R1, CCL2, CCL5, CCL22, CCL27, XCL1, CXCR1, CXCR2, CXCR 6; and pain perception PPARA* were significantly attenuated (Figure 5B). MCC950 also manifested similar effects on transcriptional level. IPA analysis demonstrated that both drugs significantly downregulated various inflammatory signals (Figure 5B and 5D). Bubble chart of in-depth IPA analysis of qPCR panels showed that rapamycin attenuated multiple signaling pathways involved in cell growth, proliferation, inflammation and fibrosis (Figure S18). These data demonstrate the potential of mTOR inhibition in regulating multiple signalling pathways involved in HS ^69^.

**Fig 5.**
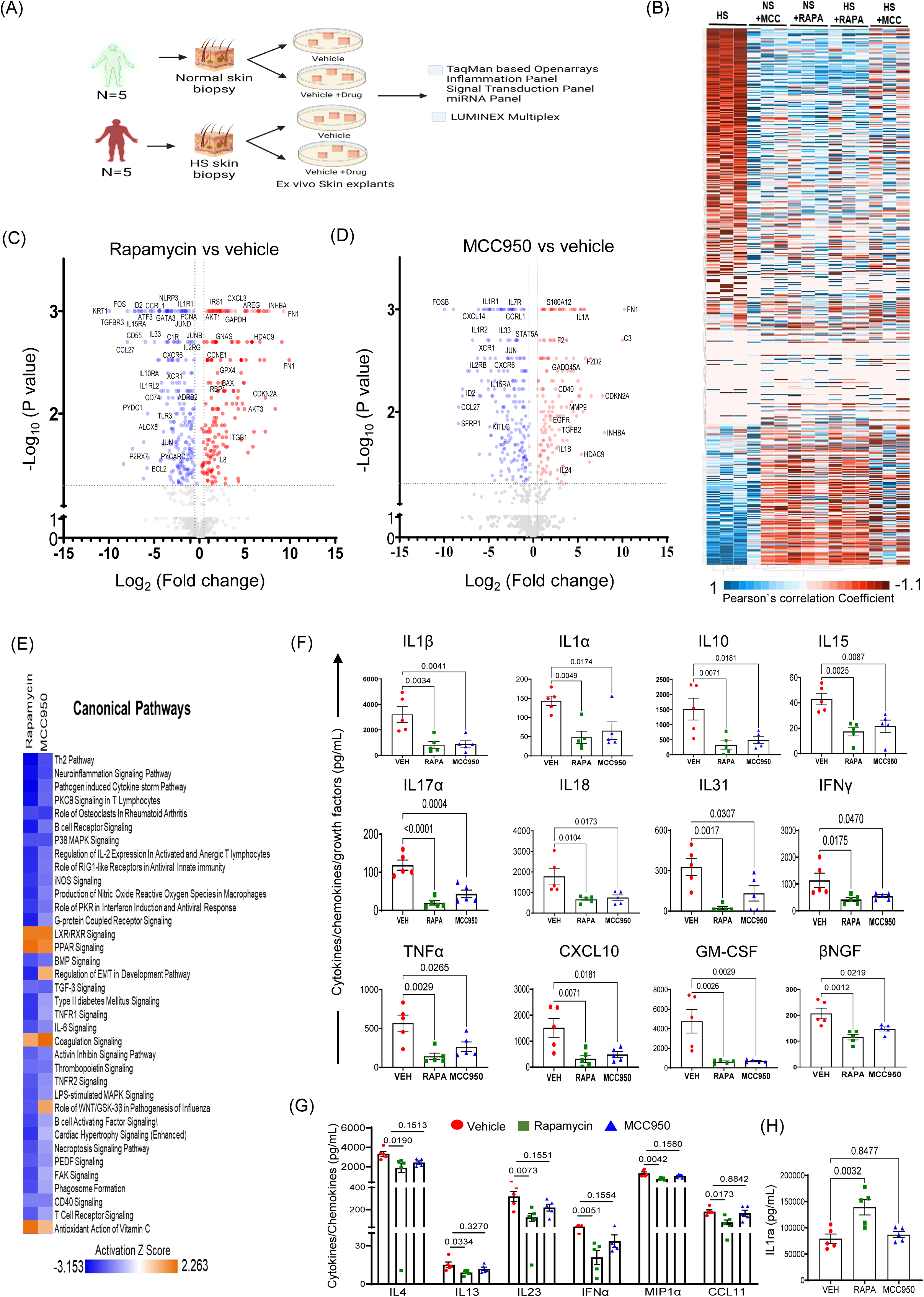
Effect of mTOR inhibitor rapamycin and NLRP3 inhibitor MCC950 on HS *ex vivo* skin cultures. **(A)** Schematic diagram illustrating drug-treatment protocol to *ex vivo* air-liquid skin explant cultures **(B)** Heatmap of genes involved in inflammation and signal transduction. Volcano plots show differentially expressed genes in rapamycin-treated **(C)** or MCC950-treated **(D)** *ex vivo* HS skin explants relative to vehicle treated HS explants (n=3/group). **(E)** Comparative analysis using IPA showing common canonical pathways in rapamycin or MCC950-treated groups. **(F)** Levels of cytokines, chemokines, and growth factors in supernatants from HS *ex vivo* skin explants cultured for 3 days. **(G)** Multiplex results demonstrate that only rapamycin significantly reduced levels cytokines/chemokines IL4, IL13, IL23, IFNα, MIP1α, and CCl11 in culture medium procured from ex vivo HS skin culture explants. **(H)** IL-1 receptor agonist is significantly induced by rapamycin. Dots in each graph represent individual HS vs NS.

To further clarify the importance of these pathways in HS, we used luminex multiplexing for assay of various proteins related to cytokines/chemokines and growth factors. Both rapamycin and MCC950 significantly (P < 0.05; n=5 each group) reduced the secretion of multiple cytokines and chemokines, including IL-1α, IL-1β, IL-10, IL-15, IL-17A, IL-18, IL-31, IFN-γ, TNF-α, CXCL10, GM-CSF, and β-NGF (Figure 5F). However, MCC950 treatment did not significantly reduce levels of IL4, IL-13, IL-23, IFN-α, MIP-1α or CCL11 (Figure 5G). The interleukin-1 receptor antagonist (IL-1Ra), which antagonizes IL-1β activity, was induced only by rapamycin (Figure 5H).

The effect of autophagy inhibitor 3-MA on HS skin was assessed to demonstrate that autophagy is instrumental in orchestrating key pathological events in this disease. For this, we treated HS skin cultures (n=5 per group) with 3MA (10mM) for three consecutive days. As expected, 3-MA synergized in augmenting inflammatory response in HS skin. Thus, we observed an exacerbated secretion of various cytokines/chemokines/growth factors in these 3 MA-treated HS cultures. The secretion of IL1β, IL5, IL7, IL13, IL15, IL17A, IL18, IL23, IL31, IFNα, IFNγ and CXCL5; and growth factors VEGFd and βNGF were elevated while anti-inflammatory cytokine IL10 was reduced as compared to baseline secretion by HS skin (Figure 6).

**Fig 6.**
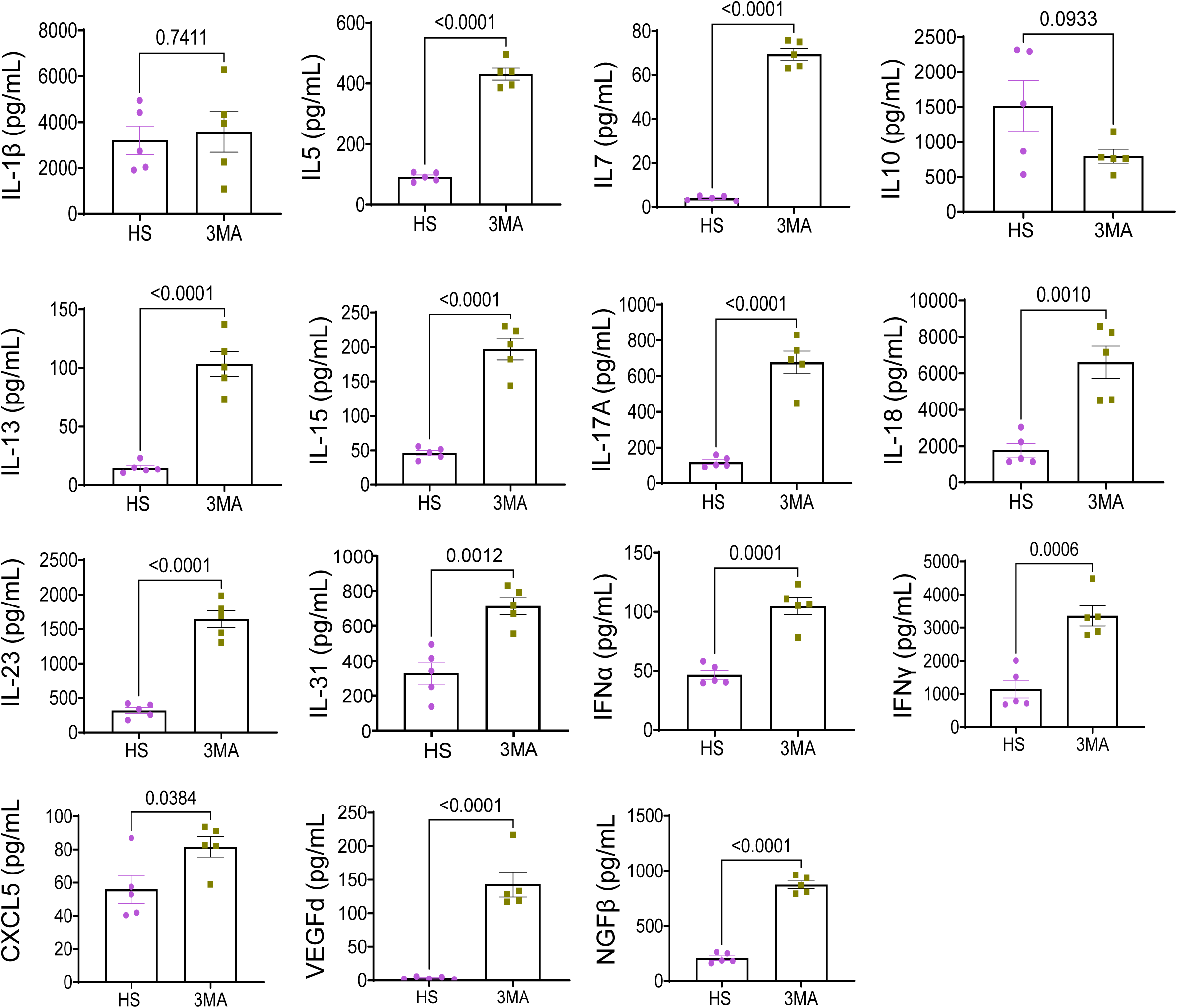
Effect autophagy inhibitor, 3 methyl-adenine (3MA) on the secretion of various chemokine/cytokines/growth factors. Effect of 3MA on the secretion of chemokine/cytokines/growth factors in culture supernatant (pooled) obtained from e*x vivo* air-liquid skin cultures at day 1 to 3. (n=5/group)

### Alterations in Epigenetic regulation in HS

To define the epigenetic mechanism of inflammatory gene signatures, we analyzed microRNA profile (n=3) of these drugs-treated HS explants. Rapamycin-treatment significantly up-regulated 5 miRs while down-modulated 39 miRs. 3MA-treatment upregulated 25 miRs and downmodulated 69 miRs whereas MCC950-treatment upregulated 26 miRs and down-modulated 48 miRs (Figure 7A). Among these, miR-99b, which is involved in mTOR inhibition was specifically and significantly upregulated by rapamycin while various miRs known for their role in autophagy inhibition were down-modulated. The examples of down-modulated miRs include: miR-200a (ATG7), miR-19a-3p and miR-19b-3p (TGF-β RII), miR-30a-5p (ATG5 and BECN1), miR-20a-5p (ATG7), miR-222 (ATG12) and miR-27a (NEDD4 and LC3II). It is recognized that rapamycin by inducing miR-642, which targets DOHH, an enzyme critical for activation of eIF5a (via hypusination) controls cell proliferation ^70^. Moreover, rapamycin-mediated reduction of miR-214 provides a mechanism for dampening NFκB-mediated inflammation by targeting MYD88 ^71^; and miR-574 by blocking JAK/STAT3 signalling ^72^ and HMGB1^73^ (Figure 7A). Synchronously, autophagy inhibitor 3MA-induced miRs having known functions in autophagy inhibition and enhancing tissue injuries. These include: miR-125a-5p (UVRAG), miR-214-3p (ATG12), miR-181A (ATG5), miR-34b-5p (BECN1), miR-222 (PTEN), miR-212 (SIRT1), miR-31 (PPP2R5A) and miR-193b-5p (OCLN). Consistently, 3MA also downmodulated many additional miRs involved in autophagy activation such as Let-7c-5p (NLRC5), has-miR-16-5p (PI3K), Let-7a-5p (BCL2L1), miR-224-5p (BCL2) (Figure 7B).

**Fig 7.**
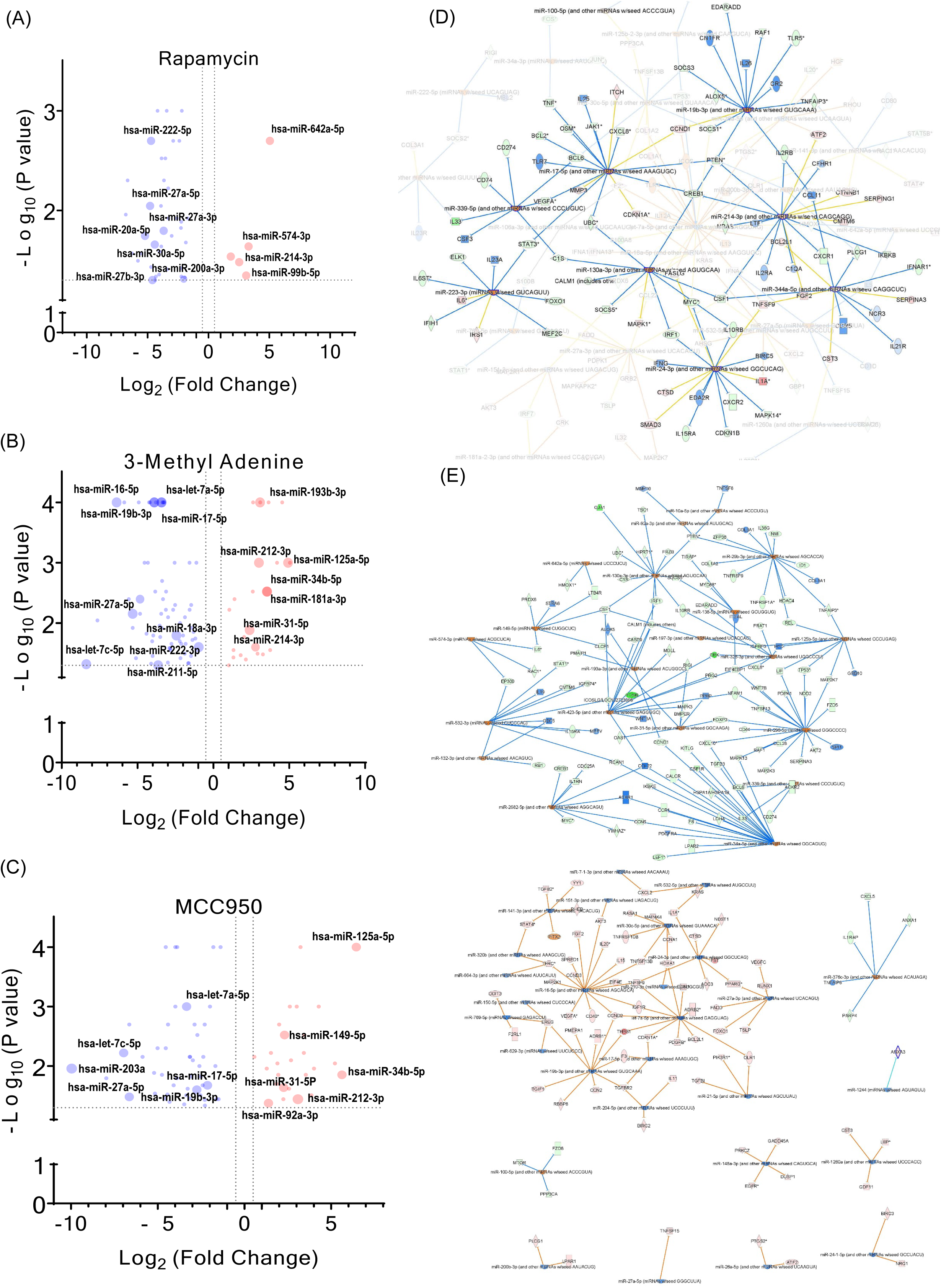
microRNA profiling and miRNA-mRNA networks of drug-treated *ex vivo* HS skin explants. Volcano plots of miRNAs following rapamycin treatment **(A)**, 3MA treatment **(B)** and MCC950 treatment groups **(C)**. IPA of miRNA-mRNA networks involved in inflammatory signaling following the treatment of rapamycin **(D)** and MCC950-regulated **(E)**.

Like rapamycin, MCC950 induced expression of miR-99b and miR-642. It also induced the expression of miRs involved in autophagy inhibition such as hsa-miR-125b-5p (BECN1), miR-138-5p (SIRT1), miR-10b-5p (PTEN), miR-339-5p (TAK1), miR-212-3p (PTEN), miR-31-5p (BECN1) and miR-34a-5p (ATG4B) (Figure 7C), providing mTOR-independent epigenetic regulation of autophagy. Further in-depth miRNA-mRNA network analysis using IPA confirmed that rapamycin induced autophagy while simultaneously reducing inflammation (Figures 7D and S19). miRNA-mRNA network analysis showed that MCC950 had influenced several independent networks, some of which work as anti-inflammatory while others work as pro-inflammatory (Figure 7E). Together these data demonstrate that autophagy is tightly epigenetically regulated in HS and provide additional layer of mechanism by which mTOR regulates HS inflammatory signaling.

## DISCUSSION

The molecular pathogenesis of HS is complex and remains incompletely defined. Our integrative multi-omics analyses reveal that HS is driven by a tightly interconnected network of metabolic, epigenetic, and cellular regulators that collectively orchestrate inflammation, pain, tissue remodeling, and disease chronicity. We uncover a distinct pathogenic signature of dysregulated metabolites—including BCAAs, BHB, NR, FBP, cadaverine and related polyamines, and multiple endocannabinoids—alongside key regulatory proteins such as ZKSCAN3, SCRN3, and mTOR. Importantly, we identify a previously unrecognized disease-associated immune population of ZKSCAN3⁺ macrophages, which function as an integrator of metabolic stress and epigenetic remodeling, orchestrating downstream inflammatory and fibrotic cascades that define the HS disease phenotype.

At the tissue level, our data reveal a coherent pathological continuum. Metabolites enriched in HS lesions contribute directly to hallmark clinical features including malodorous discharge, persistent pain, and inflammation. Notably, we observe marked increase in polyamines such as spermine, cadaverine and others. Elevated level of cadaverine and its derivatives along with elevated lysine as observed here, could sensitize pain receptors presented on afferent nerve terminals in HS patient. Consistently, we found that in KerCT keratinocytes, *in vitro* treatment with cadaverine arguments mRNA expression of multiple nociception regulatory proteins. Cadaverine also has ability to catalyse histamine toxicity by slowing down its metabolism. Histamine is known to stimulate unmyelinated c-fibres which are altered in lesional skin of HS patients ^74^. Mice receiving intradermal injection of cadaverine manifest increased scratching pattern which was shown to be regulated via histamine-4-receptors (H4Rs) and TRPV1-medited sensitization on peripheral DRG ^75^. Although polyamines can activate CD8⁺ T cells via mitochondrial engagement, our prior work demonstrates a limited contribution of CD8⁺ T cells to overall HS pathogenesis ^5^. More relevant to this study is the role of polyamines in triggering nociception via regulating endogenous activity of vanilloid (TRPV1), glutamatergic (NMDA or AMPA/kainate) receptors and acid-sensitive (ASIC) receptors ^76^. These receptors are expressed in tissues including skin, immune cells and subset of peripheral pain-sensing neurons to perceive pain signals. In this study, we also observed open reading frames and protein expression of multiple vanilloid receptors in HS skin. Significant decrease in the endocannabinoids related signalling molecules such as N-oleoyl serine, N-stearoyl serine and N-palmitoyl serine, contribute to the augmented pain pathogenesis by reducing analgesia, neuroprotection and working through PPAR-α receptor mechanism ^77,78^. Mechanistically, these phenotypes are unified by profound dysregulation of epigenetic control and autophagic flux, which emerge as central drivers of disease progression. We also confirmed that some of miRs that are known to be involved in chronic neuropathic and musculoskeletal pain regulation such as miR-21-5p, miR-29a-3p, miR-155-5p, miR-150-5p etc ^79^ are significantly altered in HS skin ^5^. In *ex vivo* HS skin explant experiment, inhibition of mTOR effectively attenuated *PPARA* gene and NGF levels, which could directly sensitize afferent nerve terminals ^80^. Topical application of mTOR inhibitor, rapamycin reduces spontaneous bleeding and pain associated skin inflammation ^81^. Consistently autophagy dysregulation is linked to the emergence of neuropathic pain in multiple diseases ^82^. Phenotypically, the convergence of aberrant protein expression and pathogenic immune–epithelial interactions underpin epithelial hyperproliferation and progressive fibrosis.

Major discovery in this manuscript is the critical role of mTOR/AMPK signaling pathways in causing severe and chronic inflammation via dysregulating autophagy. Importantly, dysregulated autophagy is observed both in the epithelial cells and immune cell compartments, which together underlie the robust NLRP3 inflammasomes driven-inflammatory responses in HS. Beyond canonical kinase signaling, mTORC1 also exerts transcriptional control over autophagy through the regulation of ZKSCAN3, a nutrient-responsive transcription factor that functions as a master repressor of autophagy and lysosomal biogenesis ^53^. Under nutrient-replete conditions, as is confirmed here by enhanced free amino acid levels in HS skin, mTORC1 activity promotes the nuclear retention and transcriptional activity of ZKSCAN3 and thereby suppressing autophagy-related gene expression. Consistent with this regulatory axis, we also identified elevated levels of metabolites such as FBP, a negative regulator of AMPK as well as increased BCAAs, both of which provide corroborative metabolic signals favouring mTORC1 activation and, by extension, ZKSCAN3-mediated repression of autophagy ^22,53,83^. In concert with these finding, several miRs, such as miR454-3p, miR-21, miR-200a and let-5d (https://mirdb.org/), which directly target tuberous sclerosis 1 (TSC1), negative regulator of mTOR ^84^ along with miRs that inhibit autophagy, such as miR-214 (ATG12), miR-376a (ATG4C and BECN), mIR-200b (ATG12), miR-200c (UBQLN1), miR-155 (ATG3) and miR-21 (Becline-1 and LC3-II), are found to be altered significantly in HS lesional skin. These data reinforce the involvement of several levels of regulatory signals in the activation of mTOR signaling in HS. Consistently, metformin, an AMPK activator and autophagy inducer was found to be beneficial in reducing HS pathogenesis in human ^85^. It may be appreciated that type 2 diabetes and obesity are intimately associated with HS ^4^ which involved dysregulated mTOR signaling and autophagy in their pathogenic cascade.

The reduced levels of NR and ketone metabolite BHB as observed in this study are also indicative of autophagy dysregulation and macrophage inflammation. Similar reduction in NR levels and associated autophagy functions is recorded in various diseases ^86^. Therefore, supplementation of nicotinamide and zinc affords protection in mild to moderate HS patients ^48^. NR significantly increases the LC3-II levels while reducing available P62 ^86^. However, in HS samples, we found significant sequestration of P62 in insoluble fraction in HS skin showing the limited availability of P62 needed for the clearance of debris in autophagosomes ^87^. The oxidative imbalance as observed in this study is likely modulator of mTOR–autophagy signaling, as oxidative stress can affect both mTOR activity and ZKSCAN3-mediated transcriptional repression of autophagy, creating a loop that amplifies autophagic impairment. These redox alterations may potentiate NLRP3 inflammasome activation and downstream inflammatory cascades, including the release of IL-1β, IL-18, TNFα, IL-17A, and IFNγ, independent of autophagy dysregulation thereby reinforcing robust sustained inflammatory signals in HS.

To further explain the role of dysregulated autophagy in keratinocytes, CosMx spatial proteomics data demonstrate that keratinocytes-specific IL18 expression in tunnel-associated epithelium and inhibited expression of LAMP1 in macrophages, NK cells and CD4^+^T cells together could lead to exacerbated inflammatory response as both keratinocytes and infiltrated macrophages express high levels of NLRP3 inflammasomes. Our data from *ex vivo* HS skin explant experiments showing that mTOR blockade not only attenuates IL-1β/IL-18 secretion, but also multiple other major cytokines/chemokines-associated with HS suggest the importance of this pathway in inflammatory HS pathogenesis.

Collectively, our findings indicate that restoring dysregulated autophagy by inhibiting aberrant mTOR activation and correcting metabolic imbalances in HS skin represents a central therapeutic opportunity. Such intervention is predicted not only to effectively suppress NLRP3 inflammasome activation and its downstream effector cytokines, IL-1β and IL-18, but also to attenuate alarmin release and broader pro-inflammatory cytokine networks, including TNFα, IL-17A, and IFNγ, all of which play well-established roles in HS pathogenesis. Together, these data position metabolic–mTOR–autophagy axis correction as a unifying strategy to mitigate chronic inflammation and disease progression in HS.

## MATERIAL AND METHODS

### Human Subject

The Institutional Review Board of the University of Alabama at Birmingham approved the study protocol (IRB-300005214) for the collection of surgically discarded skin tissues from healthy controls and HS subjects. Details regarding the use of HS and control skin samples in each experiment are provided in Table S1. Fresh tissues were processed for ex vivo skin explant culture. Portions of the tissue samples were either stored at −80 °C for molecular analyses or fixed in 10% neutral-buffered formalin and paraffin-embedded for microscopy.

### Metabolon platform

Skin tissue samples were snap-frozen and stored at −80 °C. At least 200 mg of skin tissue from either normal controls (n = 10) or HS skin samples (n = 14) were used for metabolomic and lipidomic analyses. Sample preparation, data acquisition, and data analysis were performed according to Metabolon’s standard protocols (Metabolon, Morrisville, NC, USA).

Briefly, samples were prepared using the automated MicroLab STAR® system (Hamilton Company). Proteins were precipitated with methanol under vigorous shaking for 2 min using a GenoGrinder 2000 (Glen Mills), followed by centrifugation. The resulting extract was divided into five fractions: two for analysis by separate reverse-phase (RP) UPLC-MS/MS methods with positive-ion electrospray ionization (ESI), one for RP UPLC-MS/MS with negative-ion ESI, one for HILIC UPLC-MS/MS with negative-ion ESI, and one reserved as a backup.

Samples were briefly placed on a TurboVap® (Zymark) to remove organic solvents and stored overnight under nitrogen prior to analysis. Multiple types of controls and standards were analyzed alongside experimental samples to ensure data quality. Metabolite detection was performed using ultrahigh-performance liquid chromatography–tandem mass spectrometry (UPLC-MS/MS), followed by data extraction, compound identification, curation, metabolite quantification, and normalization.

### Complex lipid platform

Lipids were extracted from samples using a methanol:dichloromethane solvent system in the presence of internal standards. The extracts were concentrated under nitrogen and reconstituted in 0.25 mL of 10 mM ammonium acetate in dichloromethane:methanol (50:50, v/v). Samples were transferred to inserts and placed in vials for infusion-based mass spectrometry analysis, performed using a Shimadzu LC system equipped with nano-PEEK tubing and a SCIEX SelexION-5500 QTRAP mass spectrometer.

Analyses were conducted in both positive- and negative-ion electrospray modes. Data acquisition on the 5500 QTRAP was performed in multiple reaction monitoring (MRM) mode, with more than 1,100 MRMs monitored. Individual lipid species were quantified by calculating the peak area ratios of target analytes to their corresponding internal standards and multiplying by the known concentration of internal standards added to each sample. Lipid class concentrations were determined by summing all molecular species within each class, and fatty acid compositions were calculated as the proportion of each lipid class contributed by individual fatty acids ^88^.

### Pathway set enrichment and principal component analyses

Pathway set enrichment analysis (PSEA) was performed using the Metabolync portal (Metabolon; https://retiredportal.metabolon.com). Pathway enrichment scores were calculated by comparing the ratio of significantly altered compounds within a given pathway to the ratio of significantly altered compounds among all named compounds detected in the study.

Principal component analysis (PCA) was conducted as an unsupervised dimensionality-reduction approach. Total variance was defined as the sum of variances of the predicted values for each principal component, and the proportion of total variance explained by each component was calculated by Metabolon (Morrisville, NC, USA). It should be noted that the Metabolon platform used in this study does not detect high-energy compounds or volatile metabolites, some of which are relevant to the biological interpretations presented here, including acetyl-CoA, di- and triphosphonucleotides, acetate, and oxaloacetate. Therefore, conclusions regarding these metabolites are inferred based on established metabolomic knowledge and pathway context rather than direct measurement. Label-free quantitative proteomic analysis was performed on fresh-frozen human skin tissue using an LC–MS/MS platform, including HS biopsies (n = 5) and matched normal skin controls (n = 5). Significantly altered proteins in HS samples were identified using standard thresholds (p ≤ 0.05 and absolute fold change ≥ 1). For phosphoproteomic analysis, protein identification and phosphorylation site assignment were performed using MS/MS spectral data analyzed with Scaffold PTM software. Phosphorylated peptides were further filtered based on statistical significance and effect size (p ≤ 0.1 and fold change ≥ 1.3). The resulting statistically significant protein abundance and post-translational modification (PTM) datasets were subsequently subjected to integrative and robust analyses using our recently developed computational pipeline ^89^.

### Proteomics and phospho-proteomics analysis

Label-free quantitative proteomic analysis was performed on fresh-frozen human tissue samples (n = 5 HS, n = 5 normal skin) using an LC–MS/MS system, comparing HS biopsies with matched normal skin controls. To identify the significantly altered proteins in HS samples, standard cutoff was used (p-value ≤ 0.05 and fold change ≥ |1|). For phospho-proteomic analysis, proteins and phosphorylation sites’ identification was performed on MS/MS spectral output with Scaffold PTM software. The modified peptides were further filtered based on p-value and fold change constraints (p-value <= 0.1 and Fold change >= 1.3). Subsequently, the two sets of statistically significant proteins (protein abundance and Protein PTM) were subjected to robust analysis using our recently devised computational pipeline^89^.

### Kinomics Analysis

Snap-frozen skin tissues from normal controls (n = 5) and HS patients (n = 5) were homogenized in MPER lysis buffer (Pierce/ThermoScientific, Cat. #78501) supplemented with Halt™ protease and phosphatase inhibitor cocktails (Protease Inhibitor Cocktail I, #P8340; Phosphatase Inhibitor Cocktail II, #P5726; and Phosphatase Inhibitor Cocktail III, #P0044; Sigma) at a 1:100 dilution for kinomics analysis. Kinomic profiling was performed using the PamStation®12 platform (PamGene International, Den Bosch, The Netherlands), a high-content phospho-peptide substrate microarray system. PamChips® were blocked with 2% bovine serum albumin (BSA) prior to sample loading. Tissue lysates (5 μg per well for protein tyrosine kinase [PTK] chips and approximately 1 μg per well for serine/threonine kinase [STK] chips) were loaded onto the PamChips® in the presence of PamGene-supplied kinase buffer, 100 μM ATP, and FITC-labeled anti-phosphotyrosine antibodies (for PTK) or anti-phosphoserine/threonine antibodies (for STK).

The assay mixtures were pumped through the PamChips® under dynamic conditions with real-time kinetic image acquisition using Evolve software (PamGene). Raw data were processed and analyzed using the BioNavigator software platform (PamGene).

### Cell culture

Ker-CT cells were cultured in KGM-Gold™ BulletKit™ medium (Lonza, 00192060) in humidified incubators at 37 °C with 5% CO₂. Cells at 60–70% confluency were treated with cadaverine at varying concentrations (10 µM–1 mM) for 24 or 72 h. Following treatment, total RNA was isolated using TRIzol reagent, and cadaverine-induced changes in signaling pathways were assessed using the Human Signal Transduction OpenArray panel (ThermoFisher Scientific, Cat. #4475389).

### ATAC-Seq

ATAC-seq datasets generated from keratinocytes isolated from HS and normal skin samples in our previous study were reanalyzed ^8^.

### CosMx sample preparation and visualization

Paraffin-embedded tissue sections were prepared for CosMx spatial profiling. Briefly, 5-µm sections of HS (n = 7) and normal (n = 3) skin tissues were mounted onto VWR Superfrost Plus Microscope Slides (Cat. #48311-703). The slides were shipped to Canopy Biosciences (Hayward, CA) for spatial profiling of immune-related proteins using the 64-plex Human Immuno-Oncology Protein Panel. Raw image processing and feature extraction were performed using the AToMx™ spatial informatics platform equipped with CosMx data analysis software v1.3 (NanoString). Multiple fields of view (FOVs) encompassing diverse pathological regions, including acanthosis, scar tissue, and sinus tracts, were analyzed. DNA and membrane stains were used to define cellular boundaries for accurate single-cell segmentation and analysis.

### *Ex Vivo* skin explants culture

An air–liquid interface platform for ex vivo skin explant culture was used as previously described ^64^. Briefly, approximately 5 × 5 mm pieces of HS (n=5) and normal skin (n=5) were placed in six-well transwell plates coated with extracellular matrix (ECM) and supplemented with KGM-Gold™ BulletKit™ (Lonza, Cat. #00192060). Cultures were maintained in humidified incubators at 37 °C under 5% CO₂.

When indicated, explants were treated with Rapamycin (2.5μM, Selleckchem), MCC950 (10μM, Selleckchem), 3-MA (10mM), or dimethyl sulfoxide (vehicle) for 3 days. Culture medium containing treatments was refreshed daily, and supernatants were collected each day for analysis of cytokines, chemokines, and growth factors. Supernatants from all three days were pooled to assess cumulative secretion. Tissues and supernatants were stored at −80 °C until further analysis.

### Quantitative real time PCR (qRT-PCR) analysis

We employed TaqMan-based real-time qPCR to evaluate mRNA expression levels of *NLRP3*, *PYCARD* and *CASP1* (Life Technologies; see Table S2). *ACTB* was used as endogenous control and relative quantification was performed using the –ΔΔCt method. Comprehensive profiling of inflammatory and cell signaling markers was conducted using predesigned gene panels, including the Human Inflammation OpenArray Panel (Cat. #4475389, ThermoFisher, USA) and Human Signal Transduction OpenArray Panel (Cat. #4475389, ThermoFisher, USA).

Total RNA from HS and normal skin tissues, as well as drug-treated or control skin explants, was isolated using TRIzol reagent (Cat. #15596018, Ambion). Two micrograms of RNA were reverse transcribed into cDNA using the SuperScript® VILO™ cDNA Synthesis Kit (Cat. #11754250, Life Technologies). Pre-amplification of cDNA was performed in a 25 µL reaction containing 12.5 µL of TaqMan PreAmp Master Mix (Cat. #4391128), 2.5 µL of custom TaqMan PreAmp primer pool (Cat. #4441856, ThermoFisher, USA), 2.5 µL of the reverse transcription product, and nuclease-free water to adjust the final volume. Thermal cycling conditions for pre-amplification included 95 °C for 10 min, followed by 12 cycles of 95 °C for 15 s and 60 °C for 4 min. Pre-amplification products were then incubated at 99.9 °C for 10 min. A 20-fold diluted pre-amplified product was used for final amplification according to the manufacturer’s instructions. OpenArray chips, pre-coated with primers for 1,183 targets, were read on a QuantStudio™ 12K Flex Real-Time PCR System (ThermoFisher Scientific, Life Technologies, Grand Island, NY).

For microRNA profiling, total RNA was isolated from drug-treated HS or untreated HS skin explants using the mirVana miRNA Isolation Kit (Cat. #AM1561). Reverse transcription was performed using the TaqMan miRNA Reverse Transcription Kit (Cat. #4366596, ThermoFisher Scientific) with Megaplex RT and PreAmp Pool Primers v3 (Cat. #4444750). miRNA profiling was performed using the TaqMan OpenArray Human MicroRNA Panel (Cat# 44701876), which contains 754 well-characterized human miRNA sequences from Sanger miRBase v14. Pre-amplification and amplification thermal cycling conditions were the same as described above. miRNA expression was measured using the 12K Flex RT-PCR system, and relative quantification was performed using the –ΔΔCt method. Data analysis was conducted using the online ExpressionSuite v1.3 software (ThermoFisher Scientific, Life Technologies, Grand Island, NY) with global gene normalization. Bioinformatics analysis of OpenArray data was performed using Ingenuity Pathway Analysis (IPA, Qiagen).

### Luminex platform

To evaluate drug responses in HS skin, multiplex analysis of cytokines, chemokines, and growth factors was performed on conditioned media from ex vivo skin cultures using the Cytokine/Chemokine/Growth Factor 45-Plex Human ProcartaPlex™ Panel 1 (Cat. #EPX450-12171-901, ThermoFisher, USA). Conditioned media were centrifuged at 10,000 rpm for 10 min and stored at −80 °C until analysis. Fifty microliters of each sample were used for multiplexing on a Luminex 200 instrument (Luminex Corporation, USA), as previously described (Kashyap et al., 2013).

A total of 45 target proteins were assessed, including: Th1/Th2 markers: GM-CSF, IFN-γ, IL-1β, IL-2, IL-4, IL-5, IL-6, IL-8, IL-12p70, IL-13, IL-18, TNF-α; Th9/Th17/Th22/Treg markers: IL-9, IL-10, IL-17A, IL-21, IL-22, IL-23, IL-27; Inflammatory cytokines: IFN-α, IL-1α, IL-1RA, IL-7, IL-15, IL-31, TNF-β; Chemokines: Eotaxin (CCL11), GROα (CXCL1), IP-10 (CXCL10), MCP-1 (CCL2), MIP-1α (CCL3), MIP-1β (CCL4), RANTES (CCL5), SDF-1α; Growth factors: BDNF, EGF, FGF-2, HGF, NGF-β, PDGF-BB, PlGF-1, SCF, VEGF-A, VEGF-D.

### High resolution Confocal Imaging

Skin sections were deparaffinized, rehydrated, and subjected to antigen retrieval using Antigen Unmasking Solution according to the manufacturer’s instructions (Vector Laboratories, Burlingame, CA). Sections were then blocked for 1 h at 37 °C in blocking buffer containing 5% normal goat serum in PBST (PBS + 0.4% Triton X-100) and incubated overnight at 4 °C with primary antibodies diluted in the same blocking solution (Table S2).

Sequential staining was performed to visualize multiple target proteins in a single specimen. After three washes with PBST (10 min each), sections were incubated with fluorescence-conjugated secondary antibodies (1:200, Invitrogen) and mounted using Vectashield Gold antifade medium containing DAPI. Stained sections were imaged on an FLUOVIEW FV3000 confocal microscope (Olympus, USA) equipped with the FV3000 Galvo scan unit and FV3IS-SW software version 2.3.2.169.

### Protein Extraction and Western Blot blotting

Skin protein lysates were prepared in ice-cold RIPA buffer containing 2 mM sodium orthovanadate, 1 mM PMSF, and 1× protease inhibitor cocktail (Santa Cruz Biotechnology). Homogenates were centrifuged at 10,000 rpm for 10 min, and supernatants were stored at −80 °C until use. P62 was analyzed using the insoluble protein fraction from these samples. Protein concentrations were determined using a BCA assay kit. Lysates were mixed with 4× sample buffer, boiled for 5 min at 95 °C, and subjected to SDS-PAGE. Proteins were transferred onto PVDF membranes, which were blocked with 5% non-fat dry milk in 1× TBST for 30 min at room temperature, followed by overnight incubation at 4 °C with primary antibodies.

After washing, membranes were incubated with HRP-conjugated secondary antibodies for 1 h at room temperature. Blots were developed using enhanced chemiluminescence (ECL, Immobilon Forte, Cat. #WBLUF0500, EMD Millipore, Burlington, MA) and imaged on the iBright FL1500 imaging system (Invitrogen). β-actin was used as a loading control. Densitometry analysis was performed using ImageJ software. A full list of primary antibodies and their dilutions is provided in Table S2.

### Statistical analysis

Differentially expressed genes (DEGs) and microRNAs (DEMs) obtained from the Inflammation, Signal Transduction, and miRNA OpenArray panels were identified using a standard cutoff of p ≤ 0.05 and |log₂ fold change| ≥ 1. Canonical pathway activation and inhibition were analyzed using Ingenuity Pathway Analysis (IPA) with thresholds of z-score ≥ |1| and Benjamini–Hochberg adjusted p ≤ 0.05.

Luminex data are presented as mean ± standard error of the mean (SEM). Statistical analyses were performed using either an unpaired Student’s t-test or one-way analysis of variance (ANOVA) followed by Sidak’s multiple comparisons test for comparisons among more than two groups.

## DATA AVAILABILITY

Requests for further information, resources, or reagents should be directed to the corresponding authors.

## ACKNOWLEDGEMENT

The authors gratefully acknowledge the technical and clinical staff who supported this work and meticulously collected and banked the samples. This work is in part supported by intramural UAB funds to M.A.

## AUTHORS CONTRIBUTIONS

Conceptualization and experimental design by M.P.K. and M.A. Majority of experiments were performed by M.P.K., and additional authors including R.S., S.H., L.J., provided technical input and assisted with experiments. Initial data visualization by M.P.K. and data evaluation by M.P.K., M.A., and C.R., Clinical evaluation of patients and tissue procurement by T.M. and C.A.E., Reagents/analytic tools contribution by M.A., Writing-original draft: M.P.K., Editing and critical comments by C.R. Writing-final draft: M.P.K, and M.A.

## References

1 Kashyap, M. P. et al. Advances in molecular pathogenesis of hidradenitis suppurativa: Dysregulated keratins and ECM signaling. Semin Cell Dev Biol 128, 120–129 (2022). 10.1016/j.semcdb.2022.01.006

2 McCarthy, S. Hidradenitis Suppurativa. Annu Rev Med 76, 69–80 (2025). 10.1146/annurev-med-051223-031234

3 Krajewski, P. K. et al. Quality-of-Life Impairment among Patients with Hidradenitis Suppurativa: A Cross-Sectional Study of 1795 Patients. Life (Basel) 11 (2021). 10.3390/life11010034

4 Cartron, A. & Driscoll, M. S. Comorbidities of hidradenitis suppurativa: A review of the literature. Int J Womens Dermatol 5, 330–334 (2019). 10.1016/j.ijwd.2019.06.026

5 Kashyap, M. P. et al. CD2 expressing innate lymphoid and T cells are critical effectors of immunopathogenesis in hidradenitis suppurativa. Proc. Natl. Acad. Sci. U. S. A. 121, e2409274121 (2024). 10.1073/pnas.2409274121

6 Jin, L. et al. Mechanism underlying follicular hyperproliferation and oncogenesis in hidradenitis suppurativa. iScience 26, 106896 (2023). 10.1016/j.isci.2023.106896

7 Adawi, W. et al. The epithelial transcriptome of hidradenitis suppurativa tunnels is more similar to cutaneous squamous cell carcinoma than to benign infundibular cysts. J Invest Dermatol (2024). 10.1016/j.jid.2024.03.051

8 Jin, L. et al. Epigenetic switch reshapes epithelial progenitor cell signatures and drives inflammatory pathogenesis in hidradenitis suppurativa. Proc. Natl. Acad. Sci. U. S. A. 120, e2315096120 (2023). 10.1073/pnas.2315096120

9 Levine, B. & Kroemer, G. Biological Functions of Autophagy Genes: A Disease Perspective. Cell 176, 11–42 (2019). 10.1016/j.cell.2018.09.048

10 Moran, B. et al. Targeting the NLRP3 inflammasome reduces inflammation in hidradenitis suppurativa skin. Br J Dermatol 189, 447–458 (2023). 10.1093/bjd/ljad184

11 Zummo, F. P. et al. Exendin-4 stimulates autophagy in pancreatic beta-cells via the RAPGEF/EPAC-Ca(2+)-PPP3/calcineurin-TFEB axis. Autophagy 18, 799–815 (2022). 10.1080/15548627.2021.1956123

12 Rames, M. M., Alavi, A. & Aghazadeh, N. GLP-1 Agonists in Patients with Hidradenitis Suppurativa: A Case Series. J Cutan Med Surg, 12034754251320045 (2025). 10.1177/12034754251320045

13 Guenin-Mace, L. et al. Dysregulation of tryptophan catabolism at the host-skin microbiota interface in hidradenitis suppurativa. JCI Insight 5 (2020). 10.1172/jci.insight.140598

14 Baumgartner, R., Forteza, M. J. & Ketelhuth, D. F. J. The interplay between cytokines and the Kynurenine pathway in inflammation and atherosclerosis. Cytokine 122, 154148 (2019). 10.1016/j.cyto.2017.09.004

15 Kostandy, B. B. The role of glutamate in neuronal ischemic injury: the role of spark in fire. Neurol Sci 33, 223–237 (2012). 10.1007/s10072-011-0828-5

16 Bennett, G. J. Update on the neurophysiology of pain transmission and modulation: focus on the NMDA-receptor. J Pain Symptom Manage 19, S2–6 (2000). 10.1016/s0885-3924(99)00120-7

17 Johnston, L., Dupuis, E., Lam, L. & Poelman, S. Understanding Hurley Stage III Hidradenitis Suppurativa Patients’ Experiences With Pain: A Cross-Sectional Analysis. J Cutan Med Surg, 12034754231188452 (2023). 10.1177/12034754231188452

18 Nishimura, M. et al. Upregulated Kynurenine Pathway Enzymes in Aortic Atherosclerotic Aneurysm: Macrophage Kynureninase Downregulates Inflammation. J Atheroscler Thromb 28, 1214–1240 (2021). 10.5551/jat.58248

19 Mariottoni, P. et al. Single-Cell RNA Sequencing Reveals Cellular and Transcriptional Changes Associated With M1 Macrophage Polarization in Hidradenitis Suppurativa. Front Med (Lausanne) 8, 665873 (2021). 10.3389/fmed.2021.665873

20 Dimitrion, P. et al. Mass cytometry uncovers a distinct peripheral immune profile and upregulated CD38 expression in patients with hidradenitis suppurativa. Cell Mol Immunol 20, 972–975 (2023). 10.1038/s41423-023-01037-6

21 Kelly, B. & O’Neill, L. A. Metabolic reprogramming in macrophages and dendritic cells in innate immunity. Cell Res 25, 771–784 (2015). 10.1038/cr.2015.68

22 Zhang, C. S. et al. Fructose-1,6-bisphosphate and aldolase mediate glucose sensing by AMPK. Nature 548, 112–116 (2017). 10.1038/nature23275

23 Mathew, R. et al. Functional role of autophagy-mediated proteome remodeling in cell survival signaling and innate immunity. Mol Cell 55, 916–930 (2014). 10.1016/j.molcel.2014.07.019

24 Poillet-Perez, L. et al. Autophagy promotes growth of tumors with high mutational burden by inhibiting a T-cell immune response. Nat Cancer 1, 923–934 (2020). 10.1038/s43018-020-00110-7

25 Petrasca, A. et al. Metformin has anti-inflammatory effects and induces immunometabolic reprogramming via multiple mechanisms in hidradenitis suppurativa. Br J Dermatol 189, 730–740 (2023). 10.1093/bjd/ljad305

26 Holecek, M. Why Are Branched-Chain Amino Acids Increased in Starvation and Diabetes? Nutrients 12 (2020). 10.3390/nu12103087

27 Wang, T. J. et al. Metabolite profiles and the risk of developing diabetes. Nat Med 17, 448–453 (2011). 10.1038/nm.2307

28 Strong, J. & Driscoll, M. S. Obesity in Hidradenitis Suppurativa: Are GLP-1 Receptor Agonists the New Frontier? Am J Clin Dermatol 26, 175–182 (2025). 10.1007/s40257-024-00911-x

29 Abu Rached, N., et al. Screening for Diabetes Mellitus in Patients with Hidradenitis Suppurativa-A Monocentric Study in Germany. Int J Mol Sci 24 (2023). 10.3390/ijms24076596

30 Mann, G., Mora, S., Madu, G. & Adegoke, O. A. J. Branched-chain Amino Acids: Catabolism in Skeletal Muscle and Implications for Muscle and Whole-body Metabolism. Front Physiol 12, 702826 (2021). 10.3389/fphys.2021.702826

31 Hawkins, M., Angelov, I., Liu, R., Barzilai, N. & Rossetti, L. The tissue concentration of UDP-N-acetylglucosamine modulates the stimulatory effect of insulin on skeletal muscle glucose uptake. J Biol Chem 272, 4889–4895 (1997). 10.1074/jbc.272.8.4889

32 Soria, L. R. et al. Beclin-1-mediated activation of autophagy improves proximal and distal urea cycle disorders. EMBO Mol Med 13, e13158 (2021). 10.15252/emmm.202013158

33 Huang, M. et al. 3-Hydroxybutyrate ameliorates sepsis-associated acute lung injury by promoting autophagy through the activation of GPR109alpha in macrophages. Biochem Pharmacol 213, 115632 (2023). 10.1016/j.bcp.2023.115632

34 Youm, Y. H. et al. The ketone metabolite beta-hydroxybutyrate blocks NLRP3 inflammasome-mediated inflammatory disease. Nat Med 21, 263–269 (2015). 10.1038/nm.3804

35 Shimazu, T. et al. Suppression of oxidative stress by beta-hydroxybutyrate, an endogenous histone deacetylase inhibitor. Science 339, 211–214 (2013). 10.1126/science.1227166

36 Sagar, N. A., Tarafdar, S., Agarwal, S., Tarafdar, A. & Sharma, S. Polyamines: Functions, Metabolism, and Role in Human Disease Management. Med Sci (Basel) 9 (2021). 10.3390/medsci9020044

37 Tang, X. et al. Ornithine decarboxylase is a target for chemoprevention of basal and squamous cell carcinomas in Ptch1+/- mice. J Clin Invest 113, 867–875 (2004). 10.1172/JCI20732

38 Silva, M. A. et al. Role of peripheral polyamines in the development of inflammatory pain. Biochem Pharmacol 82, 269–277 (2011). 10.1016/j.bcp.2011.04.015

39 Soliman, N. et al. Systematic review and meta-analysis of cannabinoids, cannabis-based medicines, and endocannabinoid system modulators tested for antinociceptive effects in animal models of injury-related or pathological persistent pain. Pain 162, S26–S44 (2021). 10.1097/j.pain.0000000000002269

40 Jacquot, F. et al. Lysophosphatidylcholine 16:0 mediates chronic joint pain associated to rheumatic diseases through acid-sensing ion channel 3. Pain 163, 1999–2013 (2022). 10.1097/j.pain.0000000000002596

41 Ren, J. et al. Elevated 18:1 lysophosphatidylcholine contributes to neuropathic pain in peripheral nerve injury. Reg Anesth Pain Med (2025). 10.1136/rapm-2024-106195

42 Hussain, A. et al. High-affinity olfactory receptor for the death-associated odor cadaverine. Proc. Natl. Acad. Sci. U. S. A. 110, 19579–19584 (2013). 10.1073/pnas.1318596110

43 Wohlfahrt, T. et al. PU.1 controls fibroblast polarization and tissue fibrosis. Nature 566, 344–349 (2019). 10.1038/s41586-019-0896-x

44 Nagaoka, Y. et al. SOX4 reversibly induces phenotypic changes by suppressing the epithelial marker genes in human keratinocytes. Mol Biol Rep 51, 116 (2024). 10.1007/s11033-023-09035-7

45 Yoshitomi, H. et al. Human Sox4 facilitates the development of CXCL13-producing helper T cells in inflammatory environments. Nat Commun 9, 3762 (2018). 10.1038/s41467-018-06187-0

46 Yoon, T. M. et al. SOX4 expression is associated with treatment failure and chemoradioresistance in oral squamous cell carcinoma. BMC Cancer 15, 888 (2015). 10.1186/s12885-015-1875-8

47 Wu, J. et al. Boosting NAD+ blunts TLR4-induced type I IFN in control and systemic lupus erythematosus monocytes. J Clin Invest 132 (2022). 10.1172/JCI139828

48 Molinelli, E. et al. Efficacy of oral zinc and nicotinamide as maintenance therapy for mild/moderate hidradenitis suppurativa: A controlled retrospective clinical study. J Am Acad Dermatol 83, 665–667 (2020). 10.1016/j.jaad.2020.04.092

49 Xie, Y., Li, J., Kang, R. & Tang, D. Interplay Between Lipid Metabolism and Autophagy. Front Cell Dev Biol 8, 431 (2020). 10.3389/fcell.2020.00431

50 Zabini, A., Zimmer, Y. & Medova, M. Beyond keratinocyte differentiation: emerging new biology of small proline-rich proteins. Trends Cell Biol 33, 5–8 (2023). 10.1016/j.tcb.2022.08.002

51 Perez-Tamayo, R. Pathology of collagen degradation. A review. Am J Pathol 92, 508–566 (1978).

52 Lue, H., Kleemann, R., Calandra, T., Roger, T. & Bernhagen, J. Macrophage migration inhibitory factor (MIF): mechanisms of action and role in disease. Microbes Infect 4, 449–460 (2002). 10.1016/s1286-4579(02)01560-5

53 Chauhan, S. et al. ZKSCAN3 is a master transcriptional repressor of autophagy. Mol Cell 50, 16–28 (2013). 10.1016/j.molcel.2013.01.024

54 He, C. Balancing nutrient and energy demand and supply via autophagy. Curr. Biol. 32, R684–R696 (2022). 10.1016/j.cub.2022.04.071

55 Ouyang, X. et al. ZKSCAN3 in severe bacterial lung infection and sepsis-induced immunosuppression. Lab Invest 101, 1467–1474 (2021). 10.1038/s41374-021-00660-z

56 Matsunaga, Y., Hashimoto, Y. & Ishiko, A. Stratum corneum levels of calprotectin proteins S100A8/A9 correlate with disease activity in psoriasis patients. J Dermatol 48, 1518–1525 (2021). 10.1111/1346-8138.16032

57 Bustin, K. A. et al. Phenelzine-based probes reveal Secernin-3 is involved in thermal nociception. Mol Cell Neurosci 125, 103842 (2023). 10.1016/j.mcn.2023.103842

58 Rudat, S. et al. RET-mediated autophagy suppression as targetable co-dependence in acute myeloid leukemia. Leukemia 32, 2189–2202 (2018). 10.1038/s41375-018-0102-4

59 Wang, B. J. et al. ErbB2 regulates autophagic flux to modulate the proteostasis of APP-CTFs in Alzheimer’s disease. Proc. Natl. Acad. Sci. U. S. A. 114, E3129–E3138 (2017). 10.1073/pnas.1618804114

60 DiPrima, M. et al. Identification of Eph receptor signaling as a regulator of autophagy and a therapeutic target in colorectal carcinoma. Mol Oncol 13, 2441–2459 (2019). 10.1002/1878-0261.12576

61 Li, Q. et al. EphA1 aggravates neuropathic pain by activating CXCR4/RhoA/ROCK2 pathway in mice. Hum Cell 36, 1416–1428 (2023). 10.1007/s13577-023-00911-9

62 Khan, A. et al. The polygenic architecture of hidradenitis suppurativa reveals signaling mechanisms that implicate epithelial remodeling. medRxiv (2025). 10.1101/2025.07.25.25332168

63 Hideshima, T. et al. Immunomodulatory drugs activate NK cells via both Zap-70 and cereblon-dependent pathways. Leukemia 35, 177–188 (2021). 10.1038/s41375-020-0809-x

64 Goliwas, K. F. et al. Ex Vivo Culture Models of Hidradenitis Suppurativa for Defining Molecular Pathogenesis and Treatment Efficacy of Novel Drugs. Inflammation 45, 1388–1401 (2022). 10.1007/s10753-022-01629-w

65 Salcedo, T. W., Kurosaki, T., Kanakaraj, P., Ravetch, J. V. & Perussia, B. Physical and functional association of p56lck with Fc gamma RIIIA (CD16) in natural killer cells. J Exp Med 177, 1475–1480 (1993). 10.1084/jem.177.5.1475

66 Wang, S. et al. S100A8/A9 in Inflammation. Front Immunol 9, 1298 (2018). 10.3389/fimmu.2018.01298

67 Navrazhina, K. et al. Epithelialized tunnels are a source of inflammation in hidradenitis suppurativa. J Allergy Clin Immunol 147, 2213–2224 (2021). 10.1016/j.jaci.2020.12.651

68 Gupta, S., Cassel, S. L., Sutterwala, F. S. & Dagvadorj, J. Regulation of the NLRP3 inflammasome by autophagy and mitophagy. Immunol Rev 329, e13410 (2025). 10.1111/imr.13410

69 Mishra, B. et al. Integrative systems biology framework discovers common gene regulatory signatures in mechanistically distinct inflammatory skin diseases. NPJ Syst Biol Appl 11, 21 (2025). 10.1038/s41540-025-00498-x

70 Epis, M. R. et al. Regulation of expression of deoxyhypusine hydroxylase (DOHH), the enzyme that catalyzes the activation of eIF5A, by miR-331-3p and miR-642-5p in prostate cancer cells. J Biol Chem 287, 35251–35259 (2012). 10.1074/jbc.M112.374686

71 Chu, Q., Sun, Y., Cui, J. & Xu, T. Inducible microRNA-214 contributes to the suppression of NF-kappaB-mediated inflammatory response via targeting myd88 gene in fish. J Biol Chem 292, 5282–5290 (2017). 10.1074/jbc.M117.777078

72 Yang, H. et al. Overexpression of miR-574-3p suppresses proliferation and induces apoptosis of chronic myeloid leukemia cells via targeting IL6/JAK/STAT3 pathway. Exp Ther Med 16, 4296–4302 (2018). 10.3892/etm.2018.6700

73 He, B. et al. MicroRNA-574-5p Attenuates Acute Respiratory Distress Syndrome by Targeting HMGB1. Am J Respir Cell Mol Biol 64, 196–207 (2021). 10.1165/rcmb.2020-0112OC

74 Lyons, D. E., Beery, J. T., Lyons, S. A. & Taylor, S. L. Cadaverine and aminoguanidine potentiate the uptake of histamine in vitro in perfused intestinal segments of rats. Toxicol Appl Pharmacol 70, 445–458 (1983). 10.1016/0041-008x(83)90162-x

75 Sun, S. Y. et al. Histamine H4 receptor and TRPV1 mediate itch induced by cadaverine, a metabolite of the microbiome. Mol Pain 20, 17448069241272149 (2024). 10.1177/17448069241272149

76 Gewehr, C. et al. Contribution of peripheral vanilloid receptor to the nociception induced by injection of spermine in mice. Pharmacol Biochem Behav 99, 775–781 (2011). 10.1016/j.pbb.2011.07.002

77 Mann, A. et al. Palmitoyl Serine: An Endogenous Neuroprotective Endocannabinoid-Like Entity After Traumatic Brain Injury. J Neuroimmune Pharmacol 10, 356–363 (2015). 10.1007/s11481-015-9595-z

78 Nakagawa-Yagi, Y. et al. Targeted lipidomics reveals changes in N-acyl serines by acute exposure to an electric field: Molecular insights into the docking of N-18:1 serine interaction with TRPV1 or PPAR-α. Integr Mol Med 6, 1–8 (2019). 10.15761/IMM.1000382

79 Sabina, S. et al. Expression and Biological Functions of miRNAs in Chronic Pain: A Review on Human Studies. Int J Mol Sci 23 (2022). 10.3390/ijms23116016

80 Kashyap, M. et al. Down-regulation of nerve growth factor expression in the bladder by antisense oligonucleotides as new treatment for overactive bladder. J Urol 190, 757–764 (2013). 10.1016/j.juro.2013.02.090

81 Hong, C. H. & Lee, C. H. Study shows an ambient-stable topical rapamycin cream is effective to treat angiofibromas in tuberous sclerosis complex. Br J Dermatol (2023). 10.1093/bjd/ljad334

82 Zheng, G., Ren, J., Shang, L. & Bao, Y. Role of autophagy in the pathogenesis and regulation of pain. Eur J Pharmacol 955, 175859 (2023). 10.1016/j.ejphar.2023.175859

83 Zhenyukh, O. et al. High concentration of branched-chain amino acids promotes oxidative stress, inflammation and migration of human peripheral blood mononuclear cells via mTORC1 activation. Free Radic Biol Med 104, 165–177 (2017). 10.1016/j.freeradbiomed.2017.01.009

84 Akkoc, Y. & Gozuacik, D. MicroRNAs as major regulators of the autophagy pathway. Biochim Biophys Acta Mol Cell Res 1867, 118662 (2020). 10.1016/j.bbamcr.2020.118662

85 Zouboulis, V. A., Zouboulis, K. C. & Zouboulis, C. C. Hidradenitis Suppurativa and Comorbid Disorder Biomarkers, Druggable Genes, New Drugs and Drug Repurposing-A Molecular Meta-Analysis. Pharmaceutics 14 (2021). 10.3390/pharmaceutics14010044

86 Kitaoka, Y. et al. Axonal Protection by Nicotinamide Riboside via SIRT1-Autophagy Pathway in TNF-Induced Optic Nerve Degeneration. Mol Neurobiol 57, 4952–4960 (2020). 10.1007/s12035-020-02063-5

87 Shi, C. S. et al. Activation of autophagy by inflammatory signals limits IL-1beta production by targeting ubiquitinated inflammasomes for destruction. Nat Immunol 13, 255–263 (2012). 10.1038/ni.2215

88 Zhang, M. et al. Adipocyte-Derived Lipids Mediate Melanoma Progression via FATP Proteins. Cancer Discov 8, 1006–1025 (2018). 10.1158/2159-8290.CD-17-1371

89 Carter, A. M. et al. Phosphoprotein-based biomarkers as predictors for cancer therapy. Proc. Natl. Acad. Sci. U. S. A. 117, 18401–18411 (2020). 10.1073/pnas.2010103117

