## Supplemental Information for "Metabolic Repression of Autophagy Drives Inflammation, Pain, and Malodor in Hidradenitis Suppurativa"

**The file includes:**

Supplementary Figures S1 -S19

Supplemental Tables S1-S2

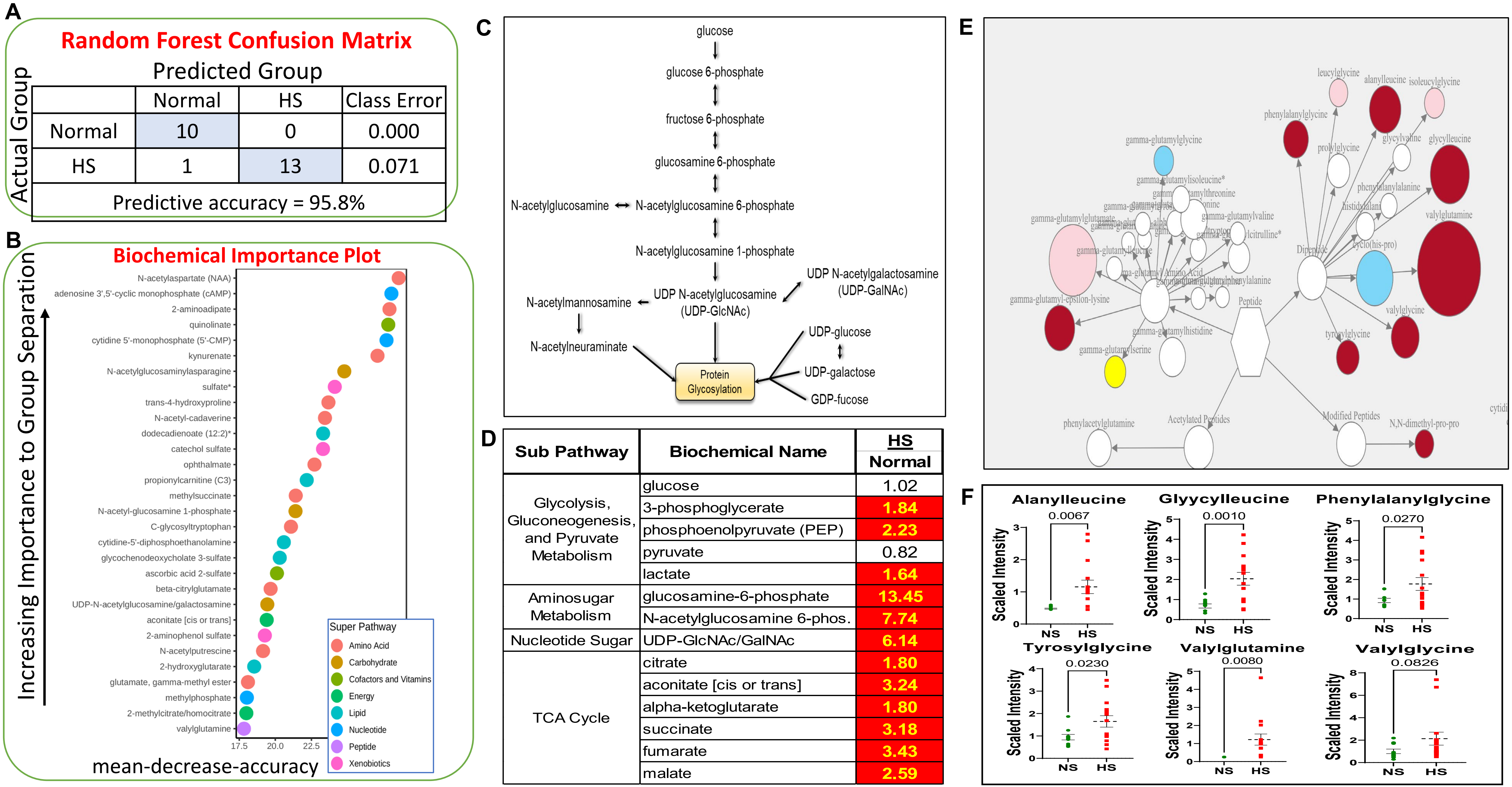

**A**

| Actual Group | Predicted Group: Normal | HS | Class Error |
|---|---|---|---|
| Normal | 10 | 0 | 0.000 |
| HS | 1 | 13 | 0.071 |
| Predictive accuracy = 95.8% | | | |

**D**

| Sub Pathway | Biochemical Name | HS / Normal |
|---|---|---|
| Glycolysis, Gluconeogenesis, and Pyruvate Metabolism | glucose | 1.02 |
| | 3-phosphoglycerate | 1.84 |
| | phosphoenolpyruvate (PEP) | 2.23 |
| | pyruvate | 0.82 |
| | lactate | 1.64 |
| Aminosugar Metabolism | glucosamine-6-phosphate | 13.45 |
| | N-acetylglucosamine 6-phos. | 7.74 |
| Nucleotide Sugar | UDP-GlcNAc/GalNAc | 6.14 |
| TCA Cycle | citrate | 1.80 |
| | aconitate [cis or trans] | 3.24 |
| | alpha-ketoglutarate | 1.80 |
| | succinate | 3.18 |
| | fumarate | 3.43 |
| | malate | 2.59 |

**Figure S1: Metabolomics of HS skin vs normal Skin.** (A) Random forest confusion matrix; (B) biochemical importance plot; (C) glucose metabolism pathway; (D) metabolite levels in HS skin; (E–F) levels of various dipeptides in HS vs normal skin samples.

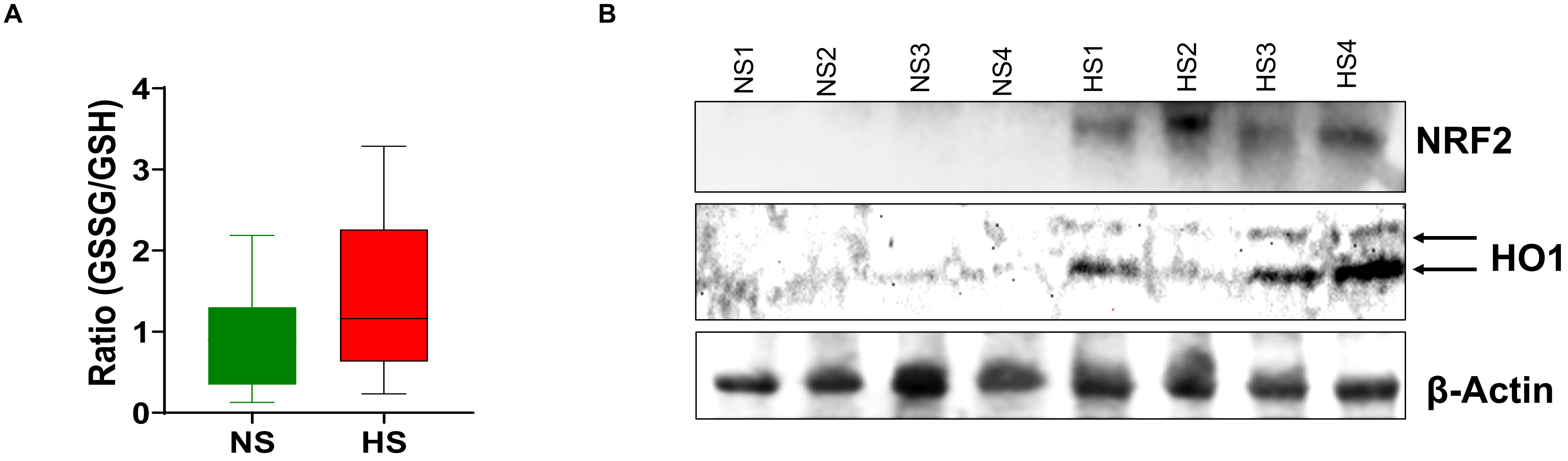

**Figure S2: NRF2-mediated antioxidant signaling is disrupted in HS.** (A) GSSG/ GSH ratio from metabolomics. (B) Western blot analysis shows increased expression of NRF2 and HO-1 in HS compared with normal skin samples.

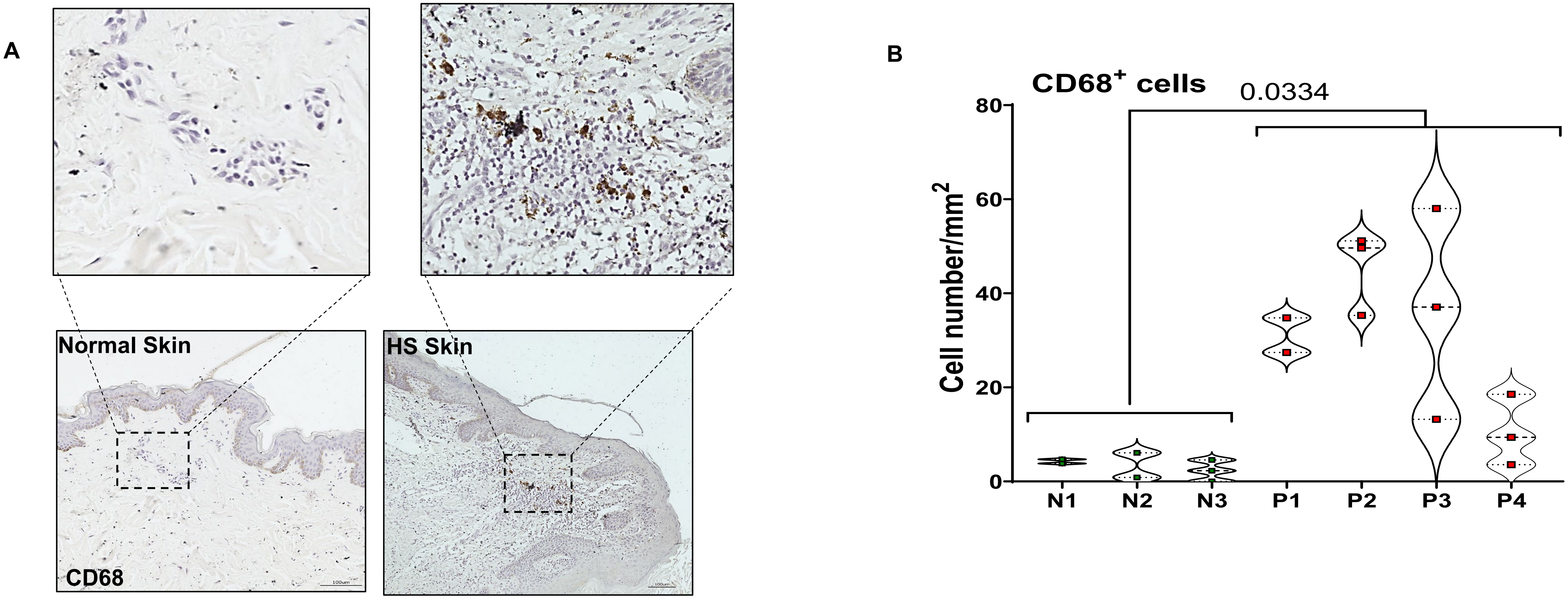

**Figure S3: Macrophage infiltration in HS skin**.(A) Immunohistochemical (IHC) analysis of normal and HS skin samples showing $CD68^+$ cells.(B) Quantification of the absolute number of $CD68^+$ cells in normal (N1–N3) and HS (P1–P4) samples. To address tissue heterogeneity and system complexity, large areas of both HS lesional and normal skin were scanned. Acquired images were stitched to generate a single high-resolution composite image of the tissue, and $CD68^+$ cells were quantified using the Macro Cell Count module of BZ-X Analyze (Keyence Corporation), which allows simultaneous analysis of several hundred (≤400) images in a single run. Depending on tissue size, each stitched image was subdivided into 3–8 regions, each containing ≤400 images, and analyzed using identical software settings. The area of each region was measured using the same software, and cell density was calculated as the number of cells per square millimeter. Statistical significance between groups was determined using an unpaired t-test with Welch's correction, with $P \leq 0.05$ considered significant.

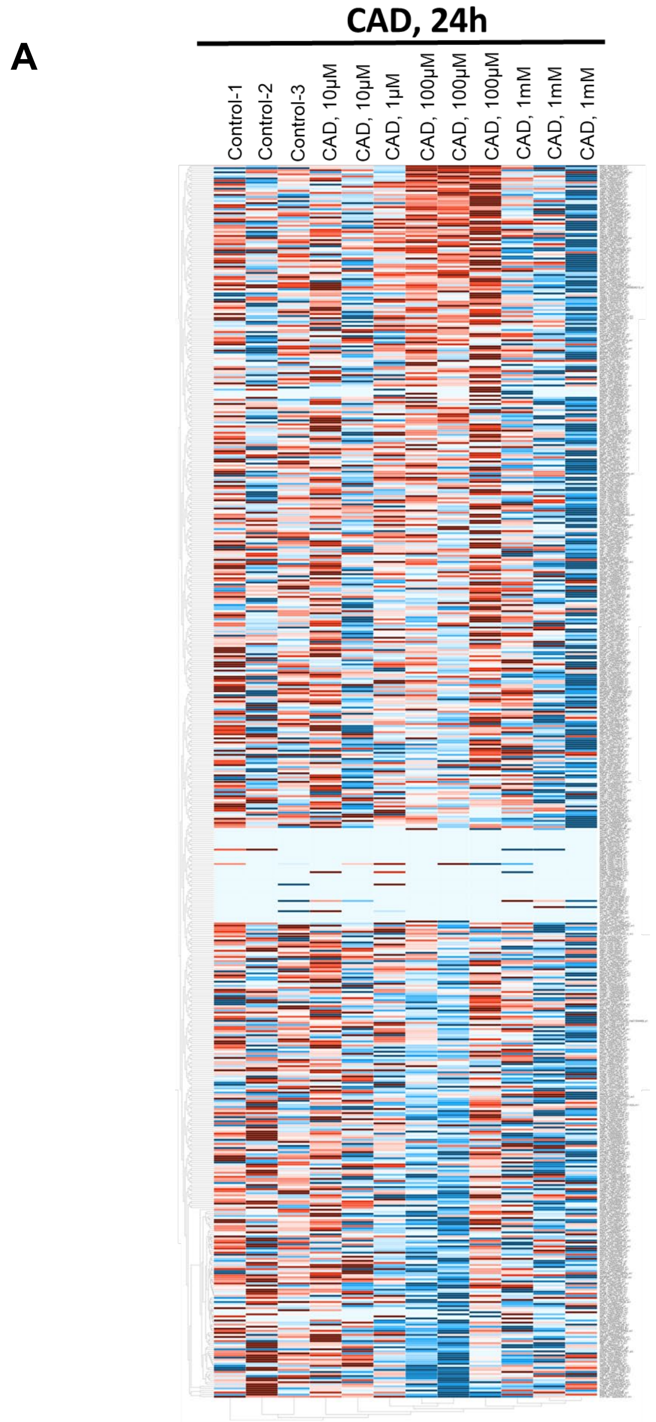

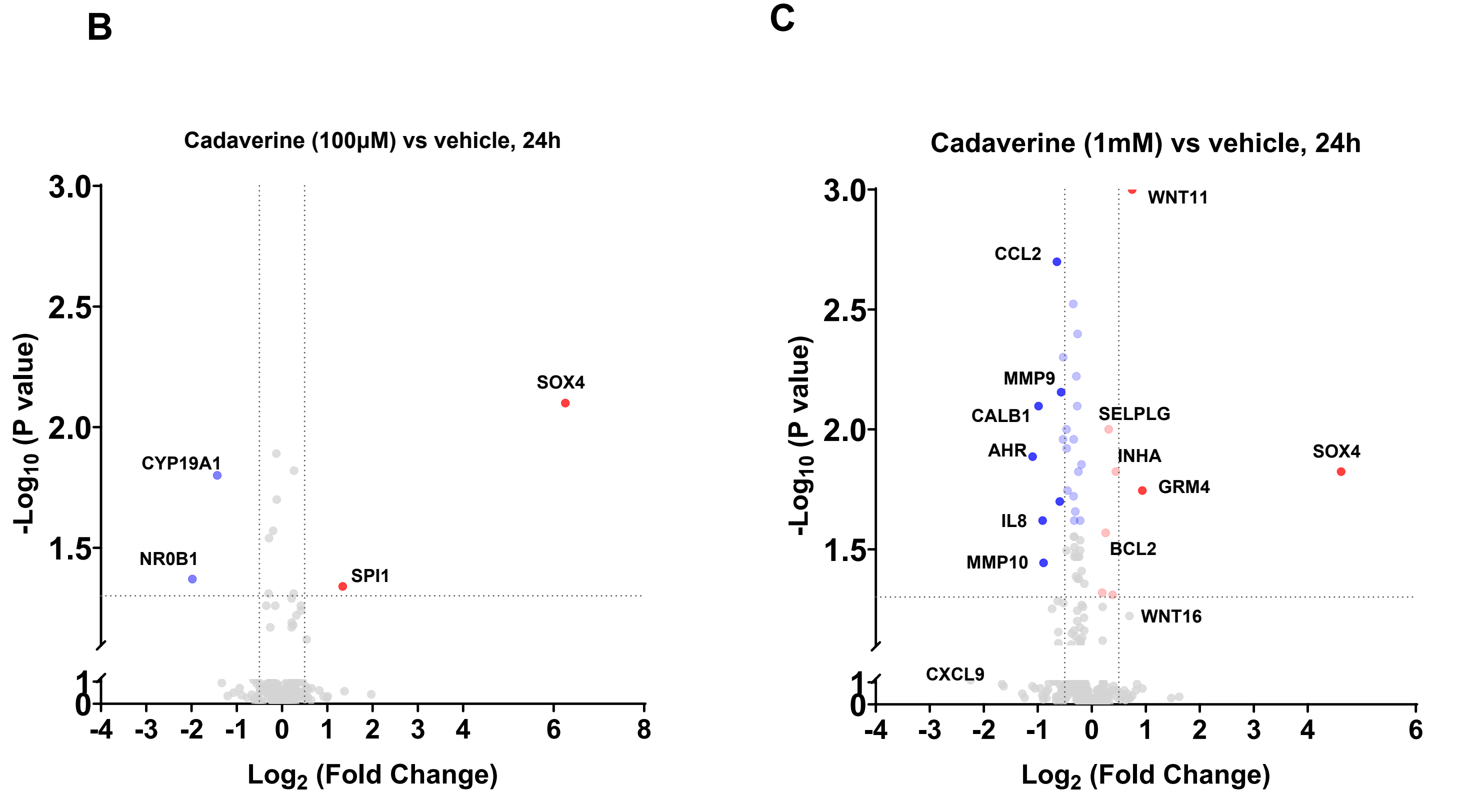

**Figure S4: Effect of cadaverine exposure on skin keratinocytes at an early time point (24 h).**

(A) Heatmaps showing genes involved in signal transduction pathways in skin keratinocytes (Ker-CT cells) exposed to increasing concentrations of cadaverine (100 μM and 1 mM; n = 3 samples/group) for 24 h.(B–C) Volcano plots depicting differentially expressed genes in Ker-CT cells following exposure to cadaverine at 1 μM (B) and 100 μM (C) for 24 h.

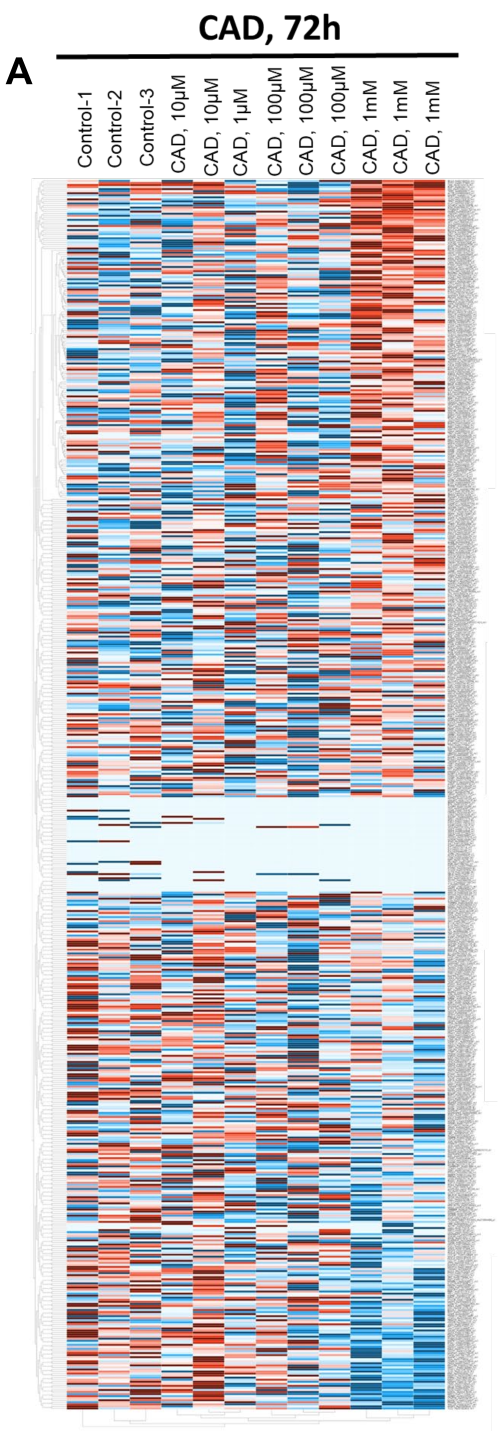

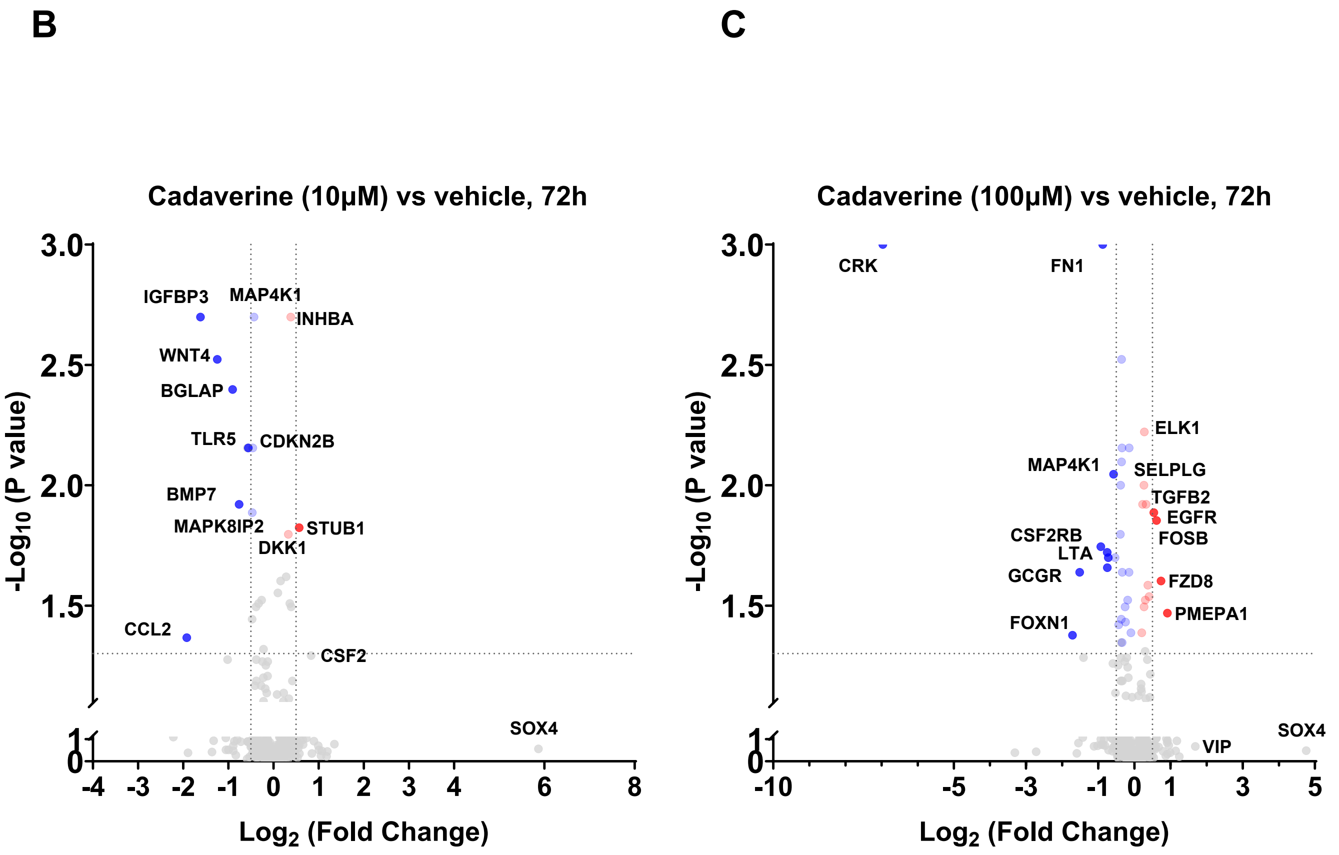

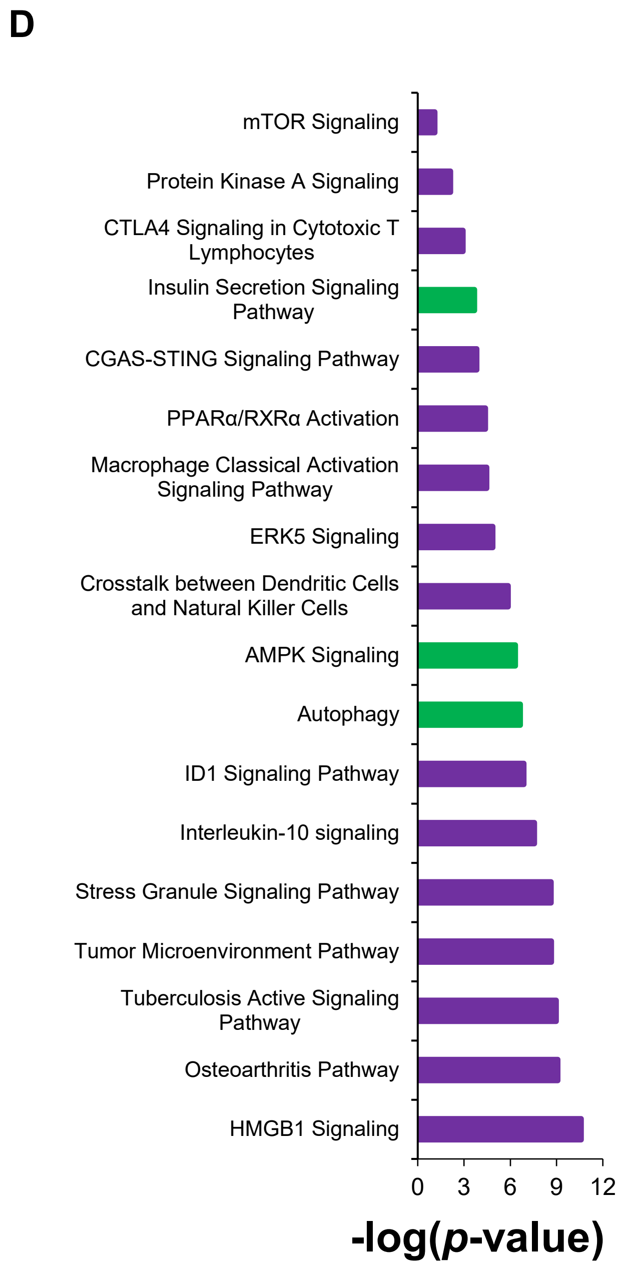

**Figure S5: Effect of cadaverine exposure on skin keratinocytes (Ker-CT cells) following prolonged treatment.**

(A) Heatmaps of genes involved in signal transduction pathways in Ker-CT cells exposed to cadaverine at 10 μM and 100 μM (n = 3 samples/group) for 72 h.(B–C) Volcano plots showing differentially expressed genes in Ker-CT cells exposed to cadaverine at 1 μM, 100 μM, and 1 mM for 48 h.(D) Ingenuity Pathway Analysis (IPA) of differentially expressed genes in Ker-CT cells exposed to cadaverine (1 mM) for 72 h. Purple bars represent activated pathways, while green bars represent downregulated pathways.

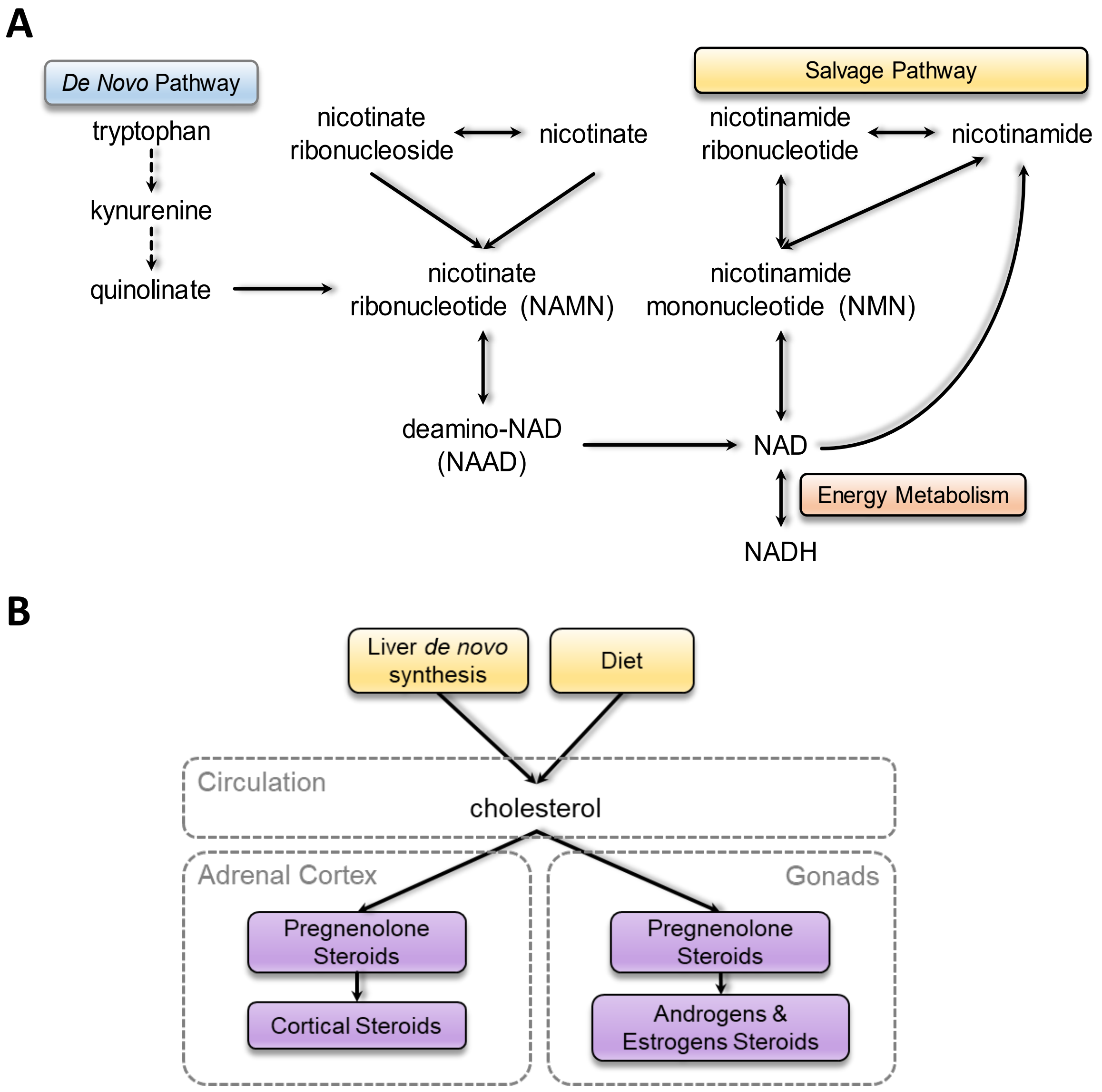

C

| Sub pathways | Symbols | HS/Normal | P value | q Value |
|---|---|---|---|---|
| Purine Metabolism, Adenine containing | Adenosine 5`-diphosphate (ADP) | 4.36 | 0.0005 | 0.0006 |
| | Adenosine 5`-monophosphate (ADP) | 7.73 | 0.0001 | 0.0002 |
| | Adenosine | 2.31 | 0.0780 | 0.0252 |
| | Adenine | 1.78 | 0.0232 | 0.0106 |
| Pyrimidine Metabolism, Uracil Containing | Uridine 3`-monophosphate (3`-UMP) | 5.14 | 0.0070 | 0.0046 |
| | uridine | 1.29 | 0.0473 | 0.0177 |
| | uracil | 7.45 | 0.0026 | 0.0021 |
| Nicotinate & nicotinamide Metabolism | Quinolinate | 241.89 | 2.51E-06 | 2.29E-05 |
| | Nicotinamide | 2.13 | 0.0018 | 0.0016 |
| | Nicotinamide ribonucleotide (NMN) | 0.47 | 0.0017 | 0.0016 |
| | Nicotinamide riboside | 0.25 | 0.0001 | 0.0003 |
| Pregnenolone Steroids | Pregnenolone Sulfate | 2.14 | 0.0764 | 0.0249 |
| | Pregnenolone disulfate* | 2.10 | 0.0209 | 0.0100 |
| Androgenic Steroids tyrosine | Androsterone glucuronide | 2.48 | 0.0156 | 0.0082 |
| | Androstenediol (3β, 17β) disulfate (1) | 2.70 | 0.0076 | 0.0048 |
| | 5α-androstane-3α, 17β-diol monosulf. | 11.93 | 0.0261 | 0.0115 |
| Primary Bile Acid Metabolism | Cholate | 8.55 | 0.0076 | 0.0048 |
| | Glycolate | 15.48 | 0.0035 | 0.0026 |
| | Taurochenodeoxycholate | 6.42 | 0.0118 | 0.0067 |

**Figure S6: Nucleotide, steroid, bile acid, and polyamine metabolism in HS skin.**
**(A–B)** Schematic diagrams illustrating nucleotide, steroid, bile acid, and polyamine metabolic pathways.
**(C)** Relative levels of metabolites involved in nucleotide, steroid, bile acid, and polyamine metabolism in HS skin relative to normal skin tissue samples.

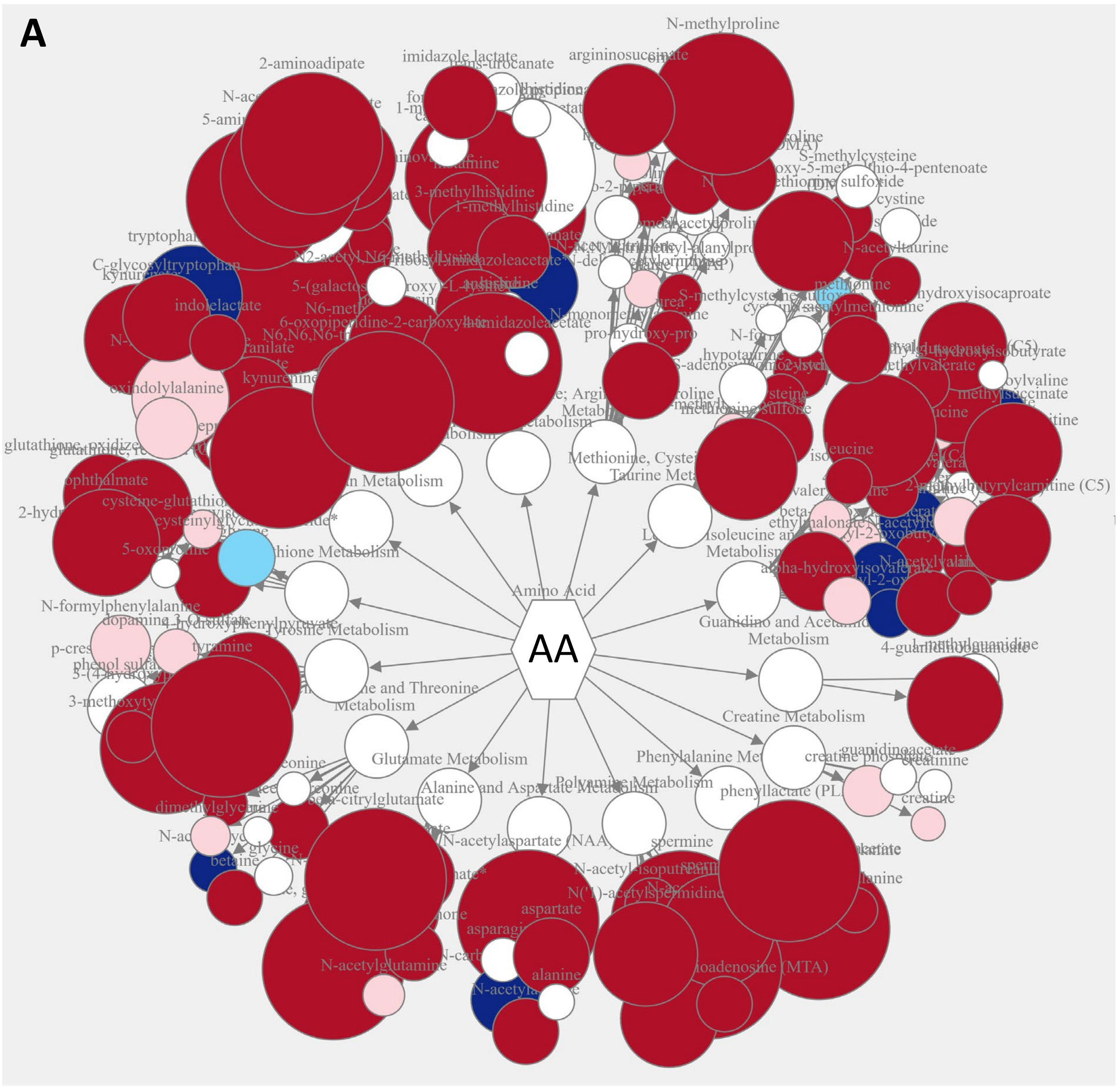

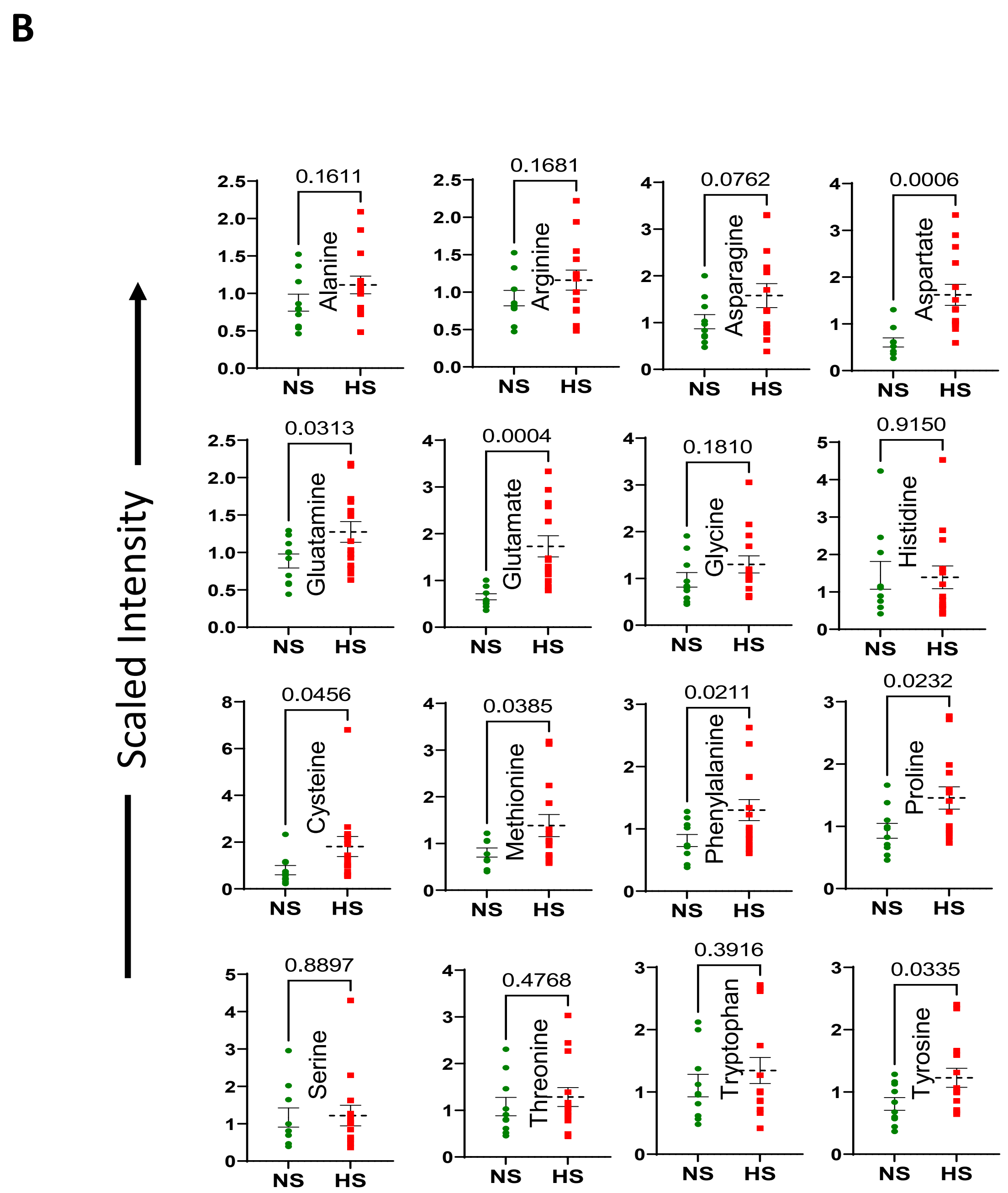

**Figure S7: Altered amino acids in HS skin.**

**(A)** Graphs showing changes in amino acid levels in HS skin relative to normal skin.

**(B)** Graphs showing the scaled intensity of various amino acids in HS and normal skin samples.

Red color indicates elevated levels, while blue indicates downregulation in HS.

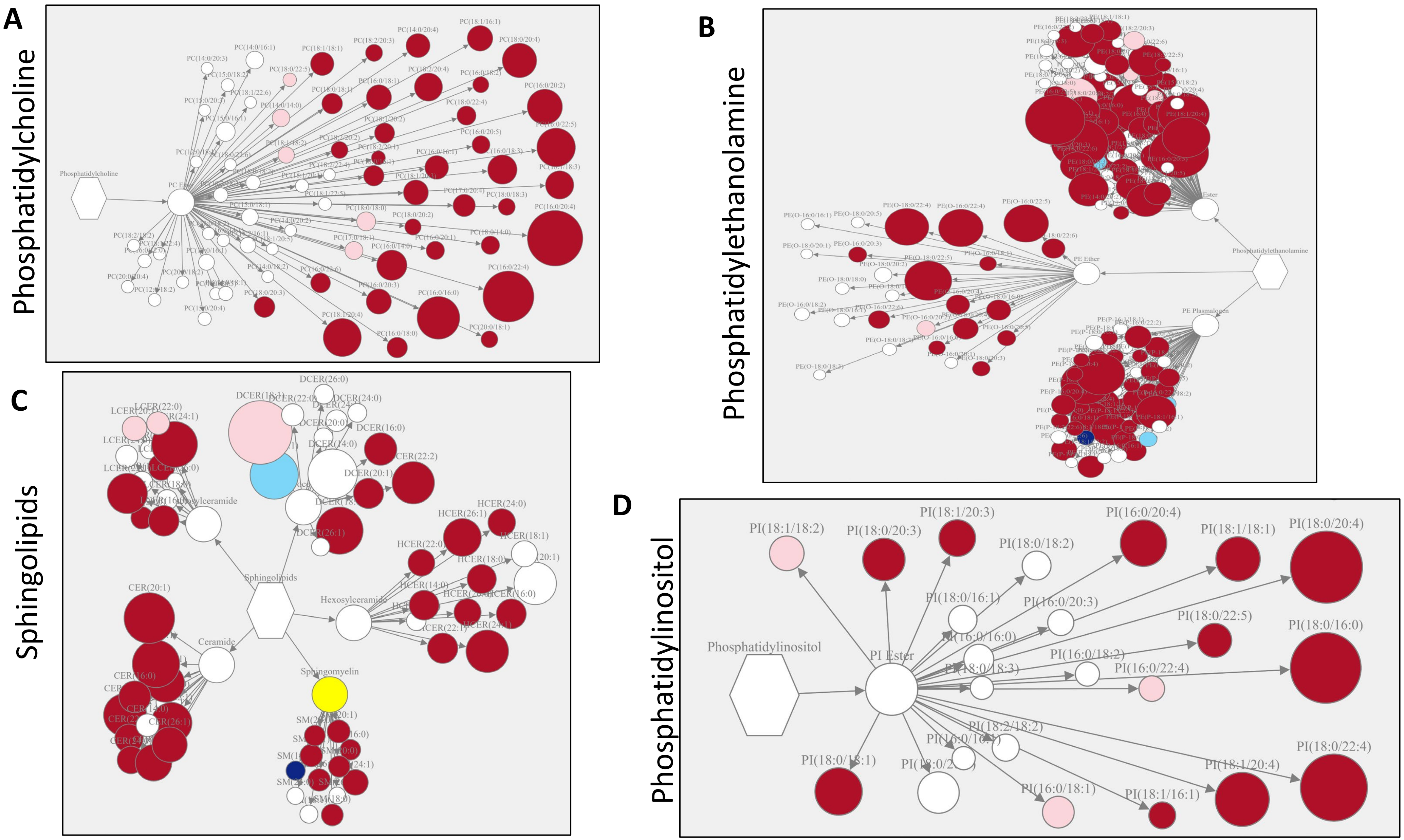

**Figure S8: Altered levels of phosphatidylcholine, phosphatidylethanolamine, sphingolipids, and phosphoinositol in HS skin. (A-D)** Diagram illustrating altered levels of phosphatidylcholine, phosphatidylethanolamine, sphingolipids, and phosphoinositol in HS skin (*n* = 14) compared with normal skin (*n* = 10). Red indicates elevated levels, blue indicates downregulation, and white indicates no change in HS. Light pink and light blue represent less pronounced alterations.

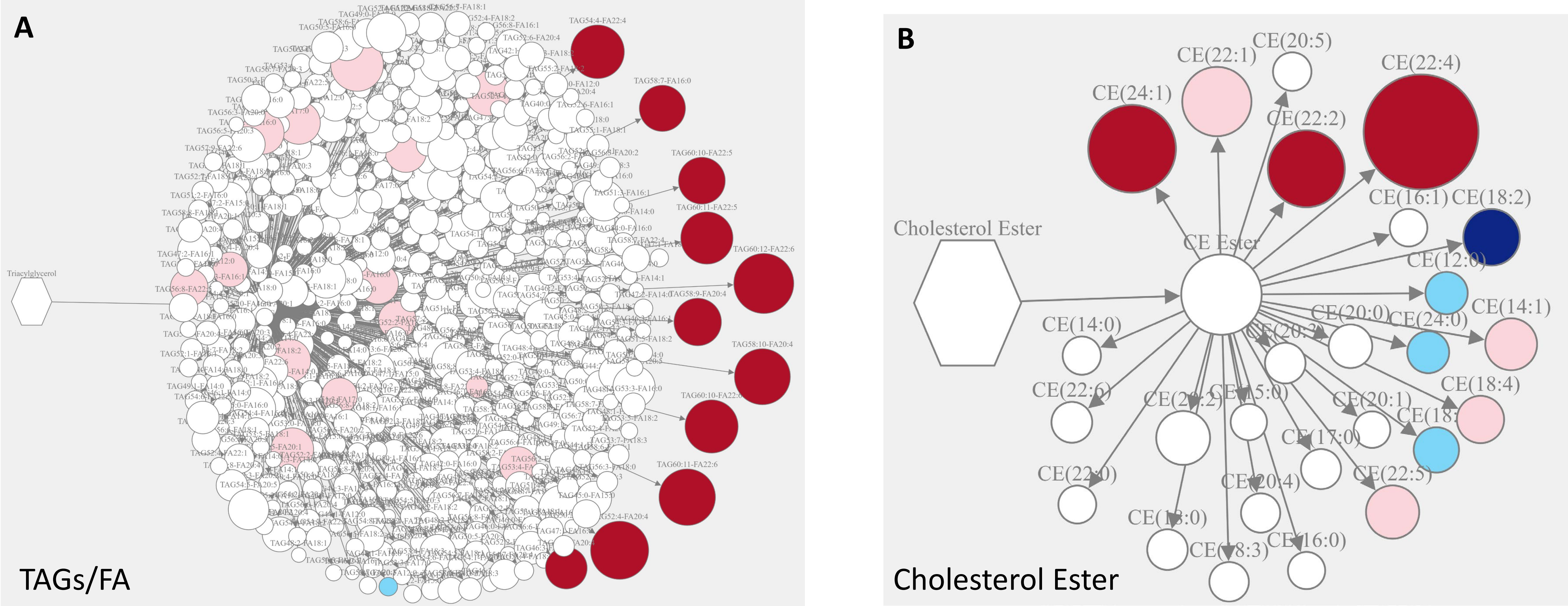

**Figure S9: Altered levels of Triglycerides (TAG) and fatty acids in HS skin. (A)** Graphs illustrating the altered TAGs/Fatty acids and **(B)** Cholesterol ester in HS skin samples relative to normal skin samples. Red indicates elevated levels, blue indicates downregulation, and white indicates no change in HS. Light pink and light blue represent less pronounced alterations.

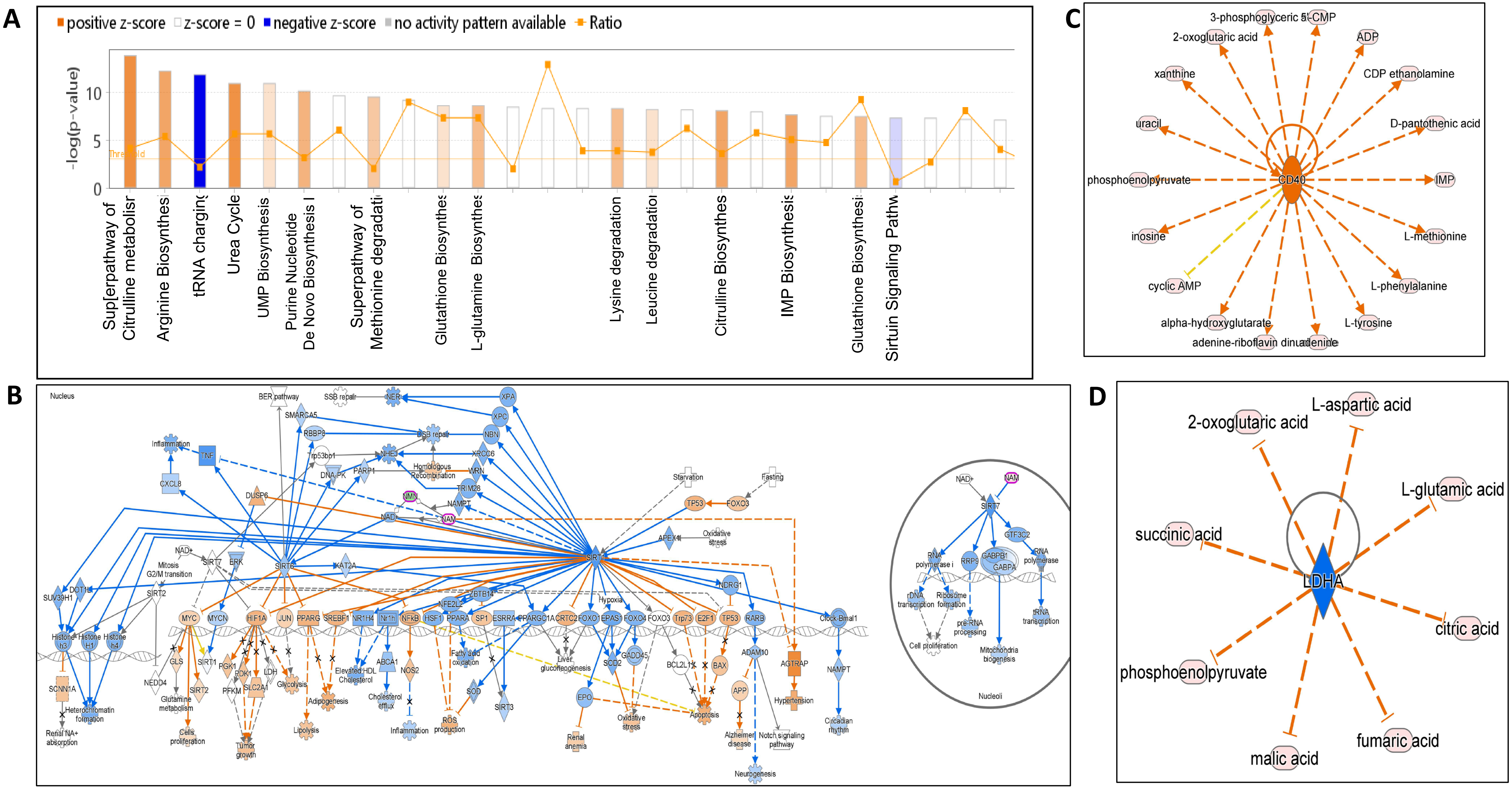

**Figure S10: IPA analysis of metabolomics data showing altered signaling in HS skin.**
**(A)** Altered pathways. **(B)** Signaling diagrams showing involvement of various genes predicted by altered metabolites. **(C–D)** Predicted upstream regulators in HS.

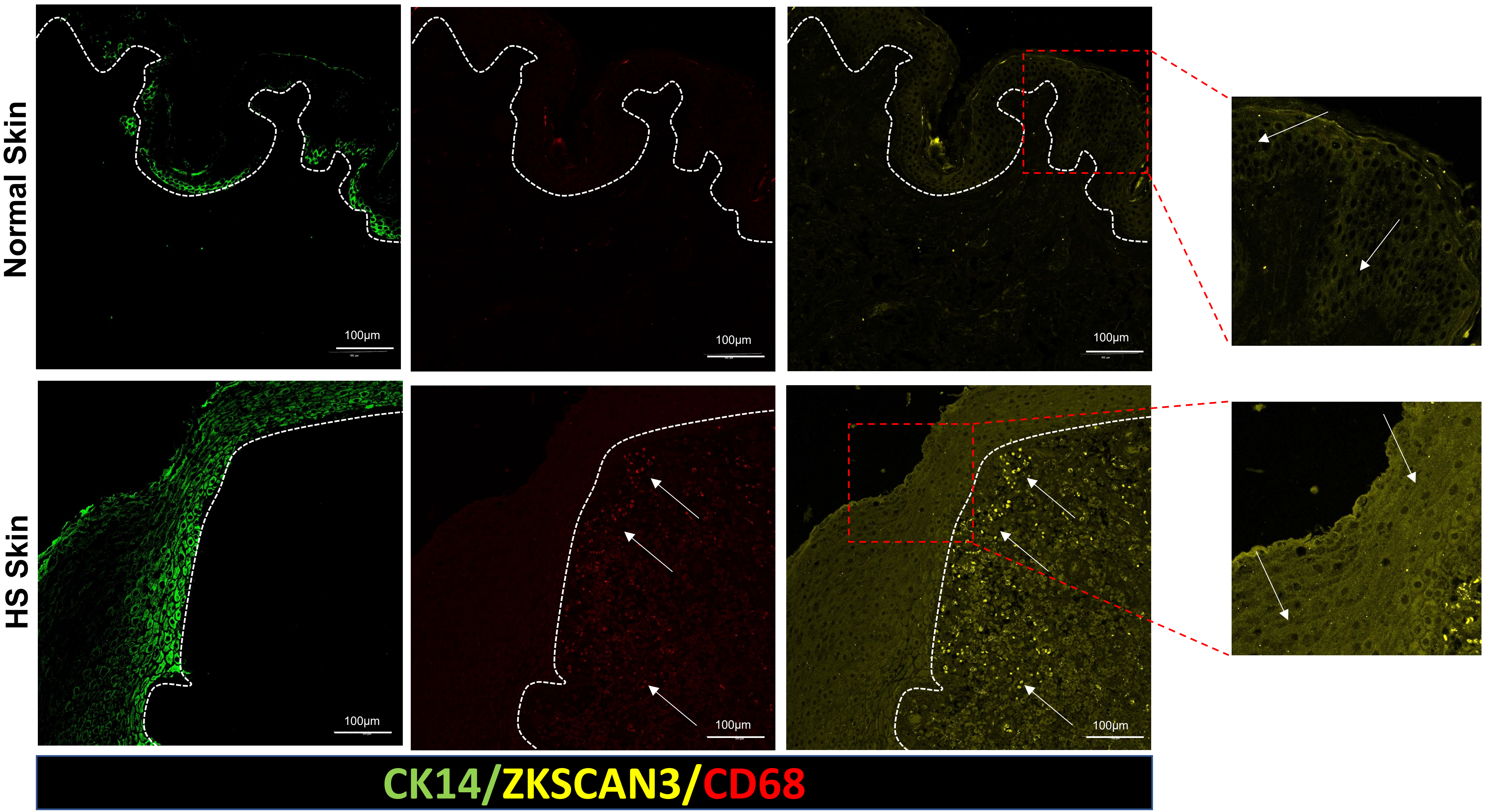

**Figure S11: ZKSCAN3 in HS skin.** Immunofluorescence micrographs showing CK14 (epidermis; green), CD68 (macrophages; red), and ZKSCAN3 (yellow) in normal and HS skin (epidermal and hypodermal layers).

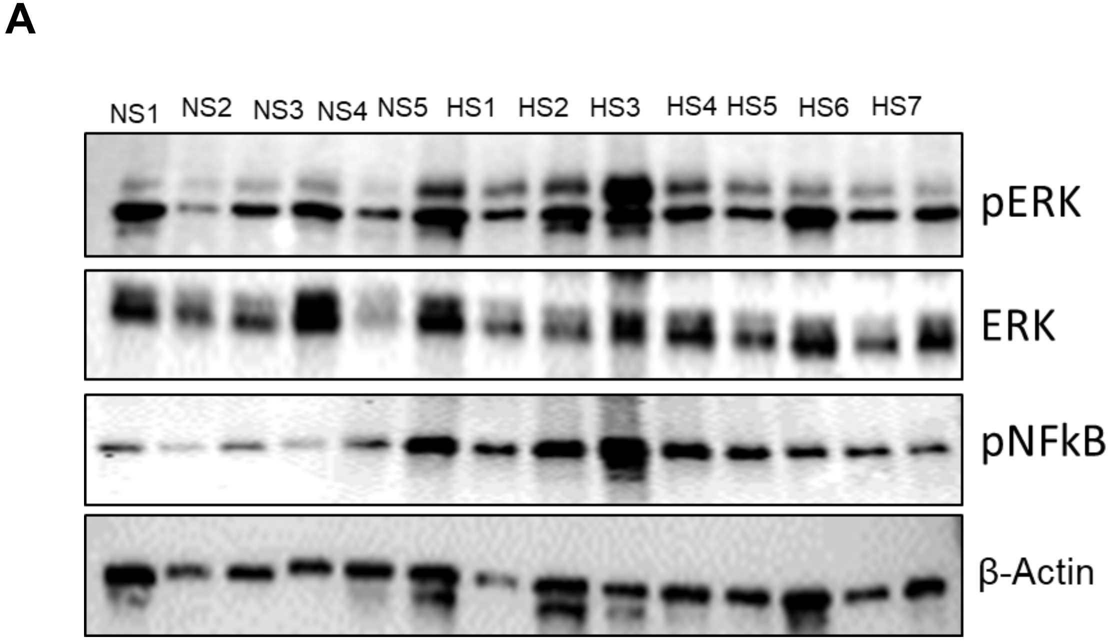

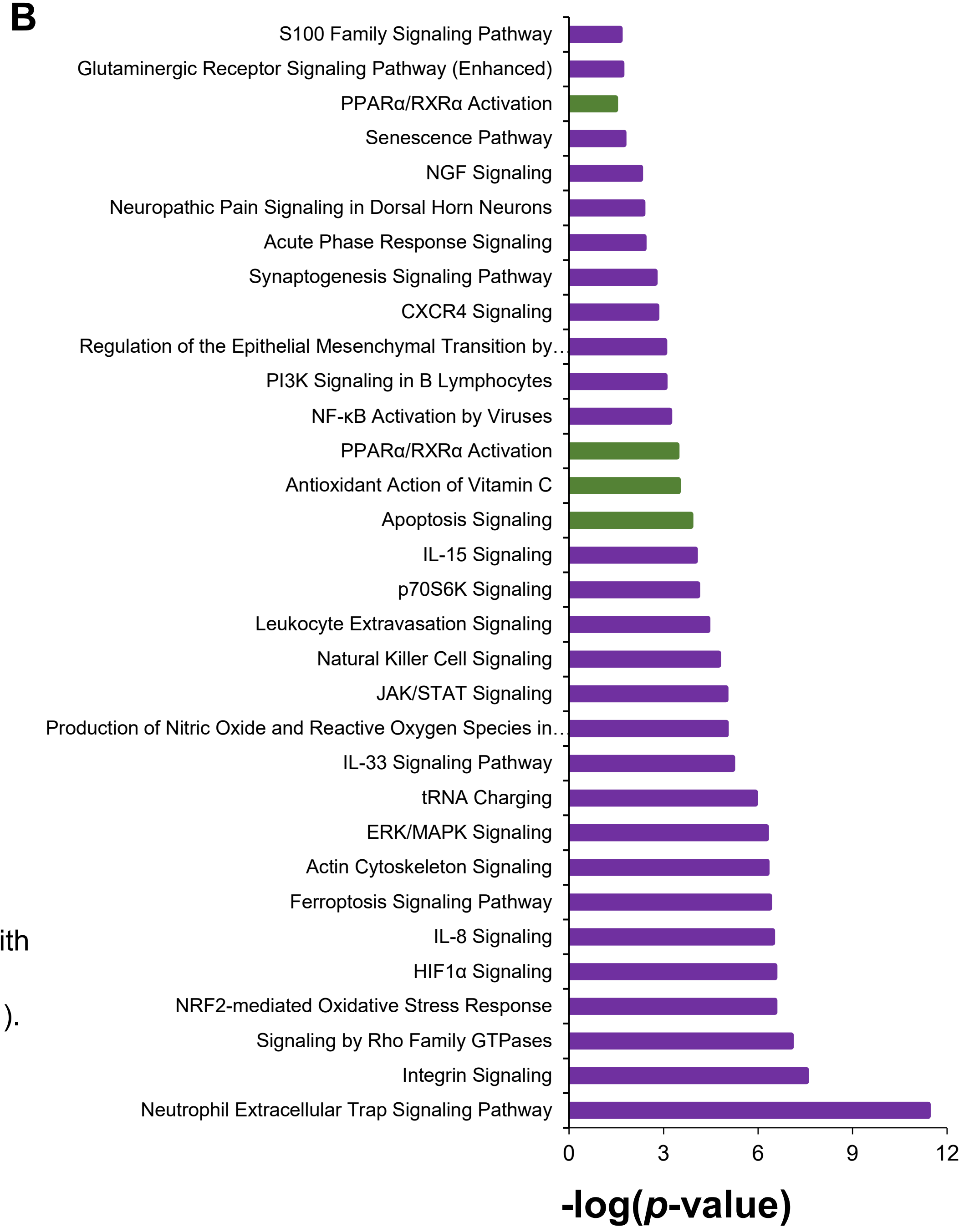

**Figure S12: Altered proteome in HS.**

**(A)** Western blot showing increased pERK and pNFKB1 in skin samples from HS patients compared with normal subjects.

**(B)** IPA analysis of proteomics data showing altered signaling in HS skin (*log* BH $p \leq 0.05$; |z-score| > 1). Purple bars represent activated pathways, while green bars represent downregulated pathways.

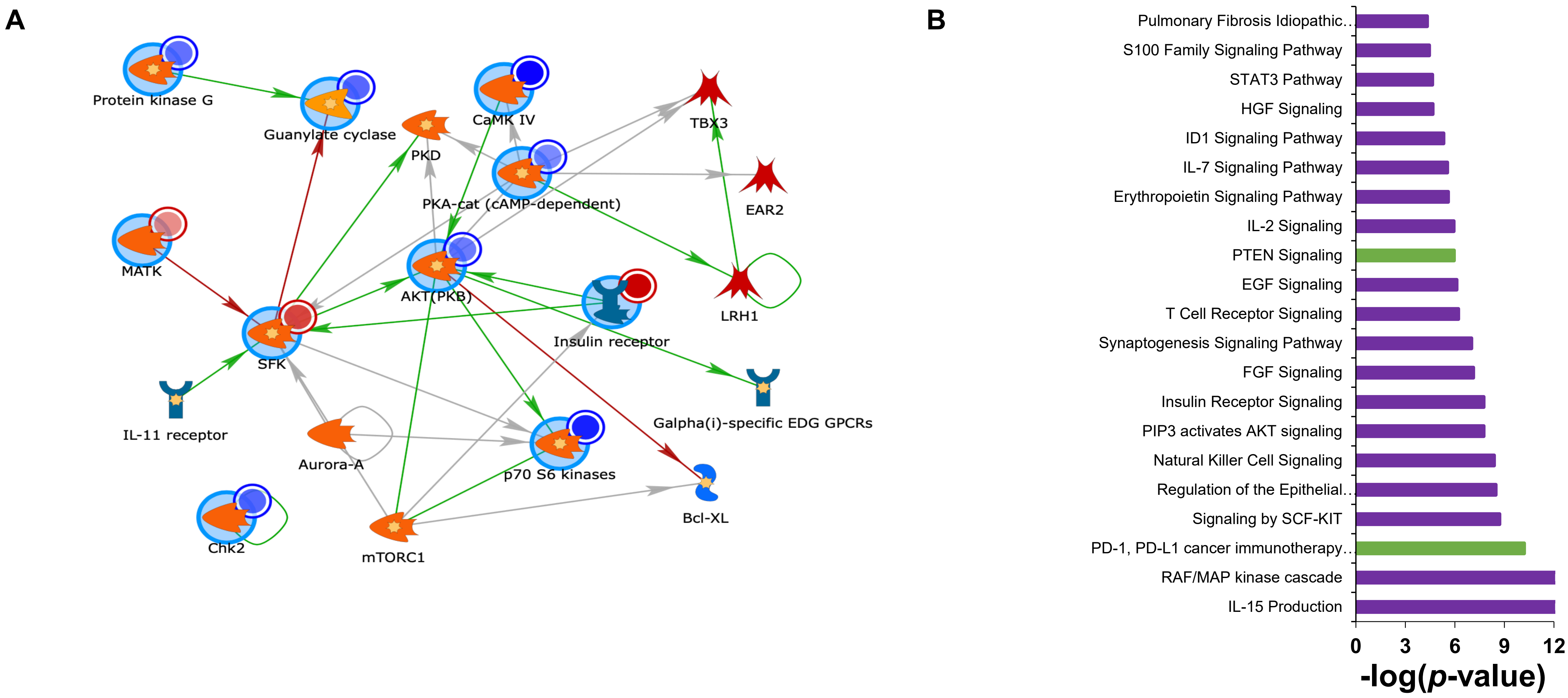

**Figure S13: Altered kinome in HS skin.**
**(A)** Network mapping of combined molecular profiling in HS. Altered kinases were uploaded to GeneGo MetaCore, and the network was modeled as an IKK-β–centric network (AutoExpand, *n* < 75). Input kinases are shown as large blue circles, with smaller circles in the top right indicating changes (red = increased in HS, blue = decreased). Arrowheads denote the direction of interaction, and line colors indicate interaction type (green = positive, red = inhibitory, grey = complex).
**(B)** IPA analysis of kinome data highlighting involvement of NK cell signaling and IL-15 production. Purple bars represent activated pathways, while green bars represent downregulated pathways.

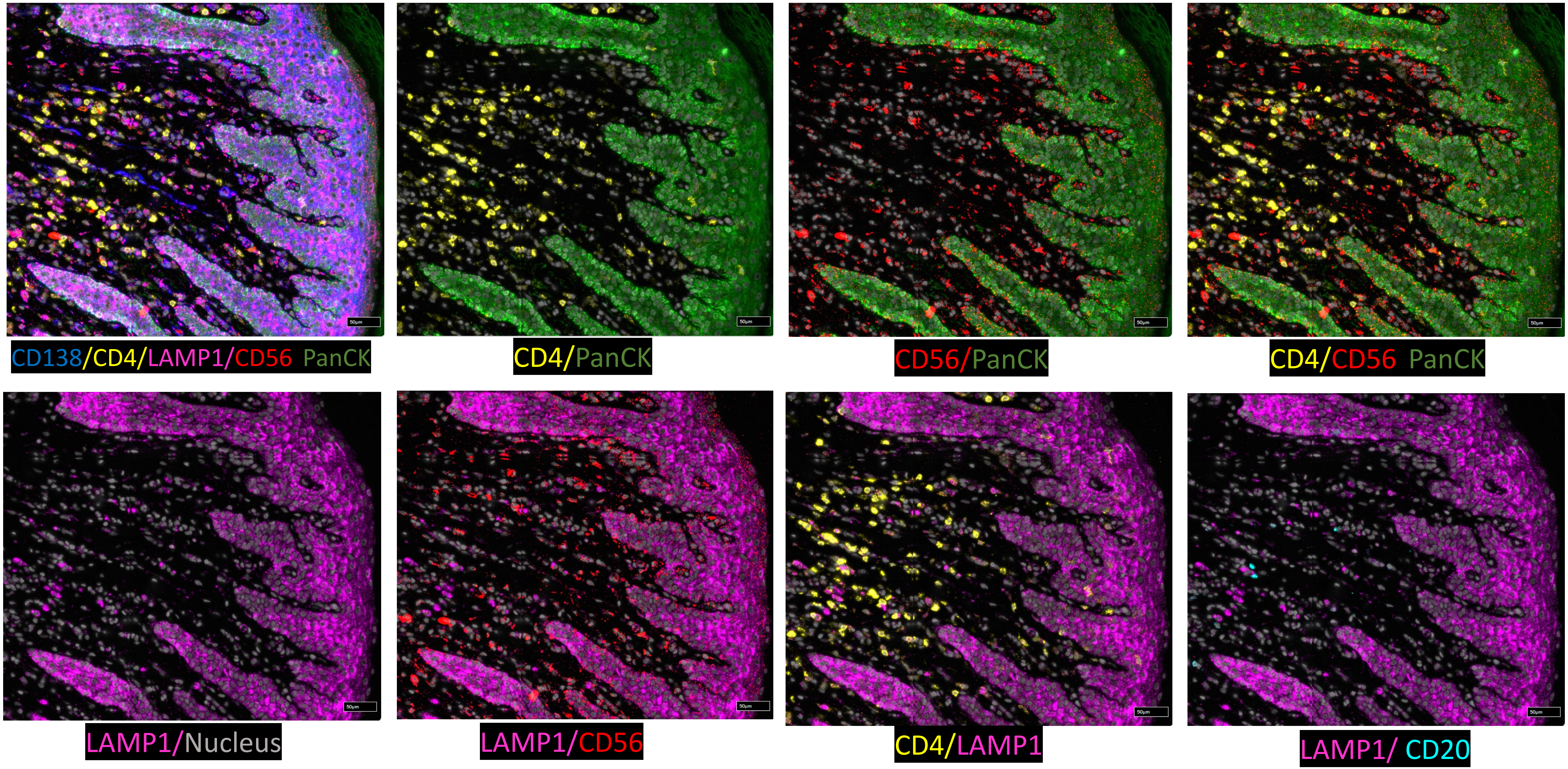

**Figure S14:** CosMx protein multiplexing images of HS skin revealing a hyperinflammatory immune environment.

A

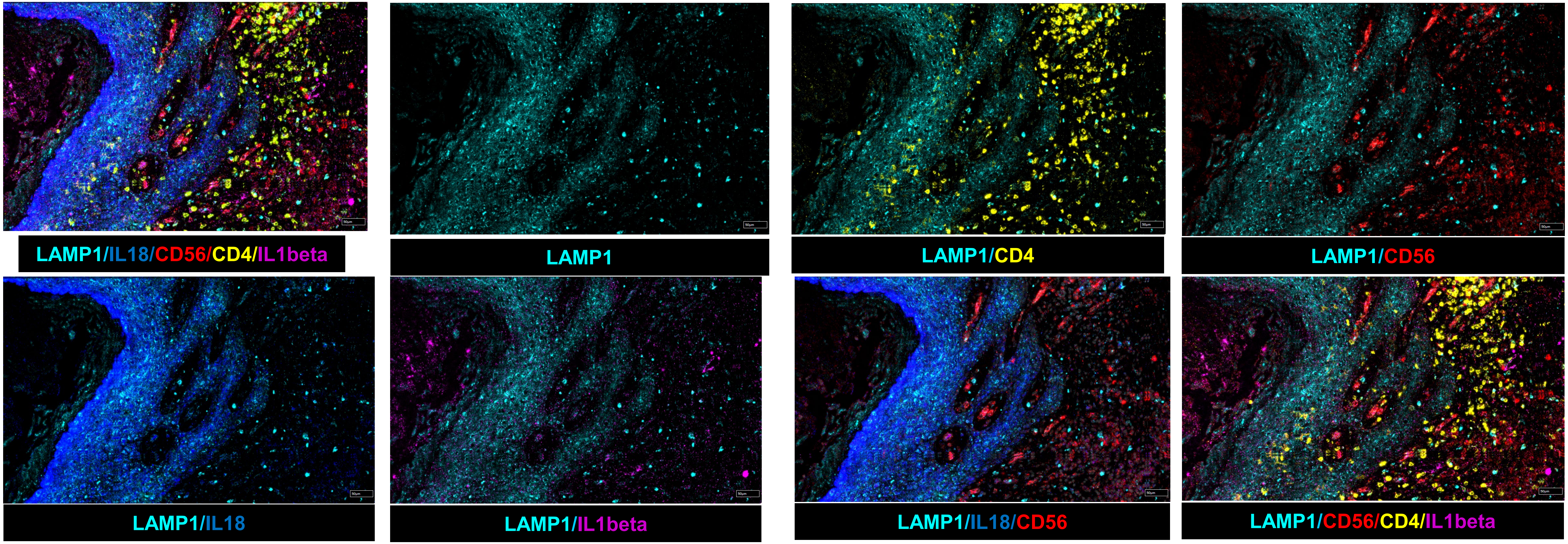

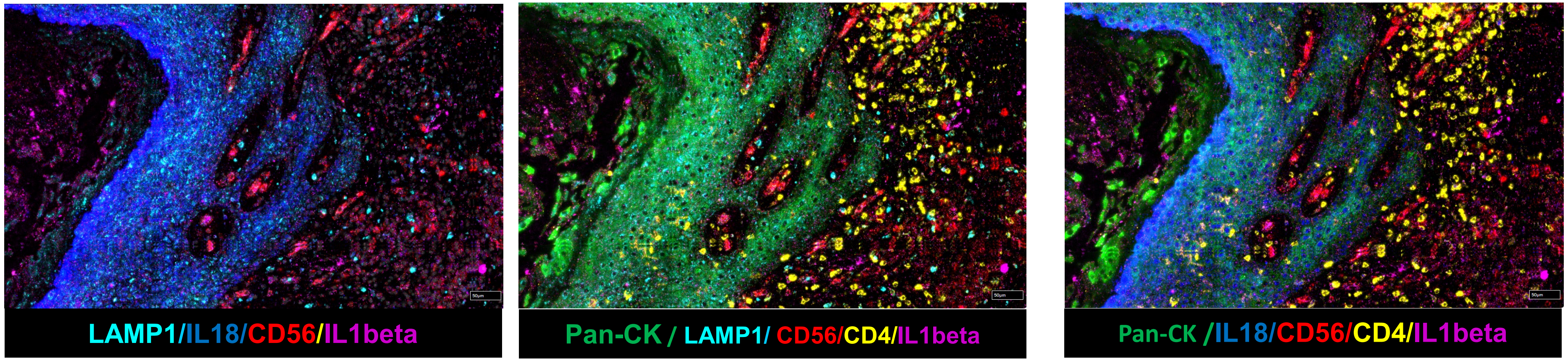

B

| cluster | Genes | avg_log2FC | p_val | p_val_adj |
|---|---|---|---|---|
| CD4TCells | MAP1LC3B | -0.80092687 | 7.24E-55 | 1.84E-50 |
| BCells_pDC | MAP1LC3B.1 | -0.94453848 | 3.73E-78 | 9.45E-74 |
| CD8T_NKTCells | MAP1LC3B.2 | -0.3720498 | 1.65E-04 | 1 |
| CD8TCells | MAP1LC3B.3 | -0.65261524 | 7.58E-16 | 1.92E-11 |
| TNaive | MAP1LC3B.6 | -1.3539463 | 5.41E-163 | 1.37E-158 |

**Figure S15:** CosMx protein multiplexing images of HS skin revealing LAMP1 on Sinus tracts including immune cells. **(B)** Down-modulated mRNA levels of LC3B in various immune cell populations from our previous scRNA Seq study (Kashyap et al, PNAS).

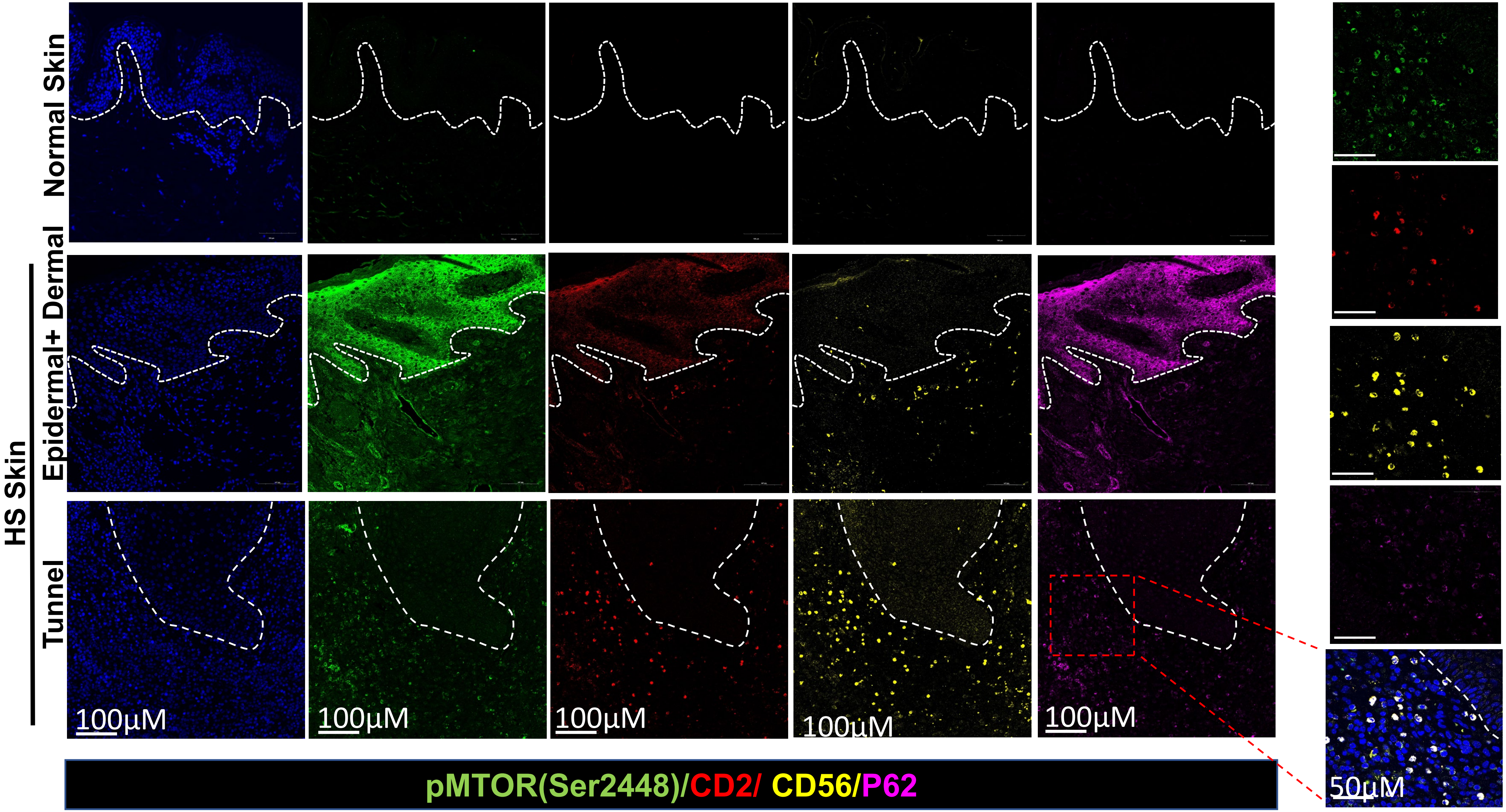

**Figure S16. mTOR signaling in HS skin.** Immunofluorescence micrographs showing p-mTOR (der2485) in CD2$^+$CD56$^+$ (NK) cells and the epidermis of normal and HS skin (epidermal/hypodermal layers, Tunnel region). Resolution: 4096 × 4096; magnification: ×20; scale bars: 100 µm.

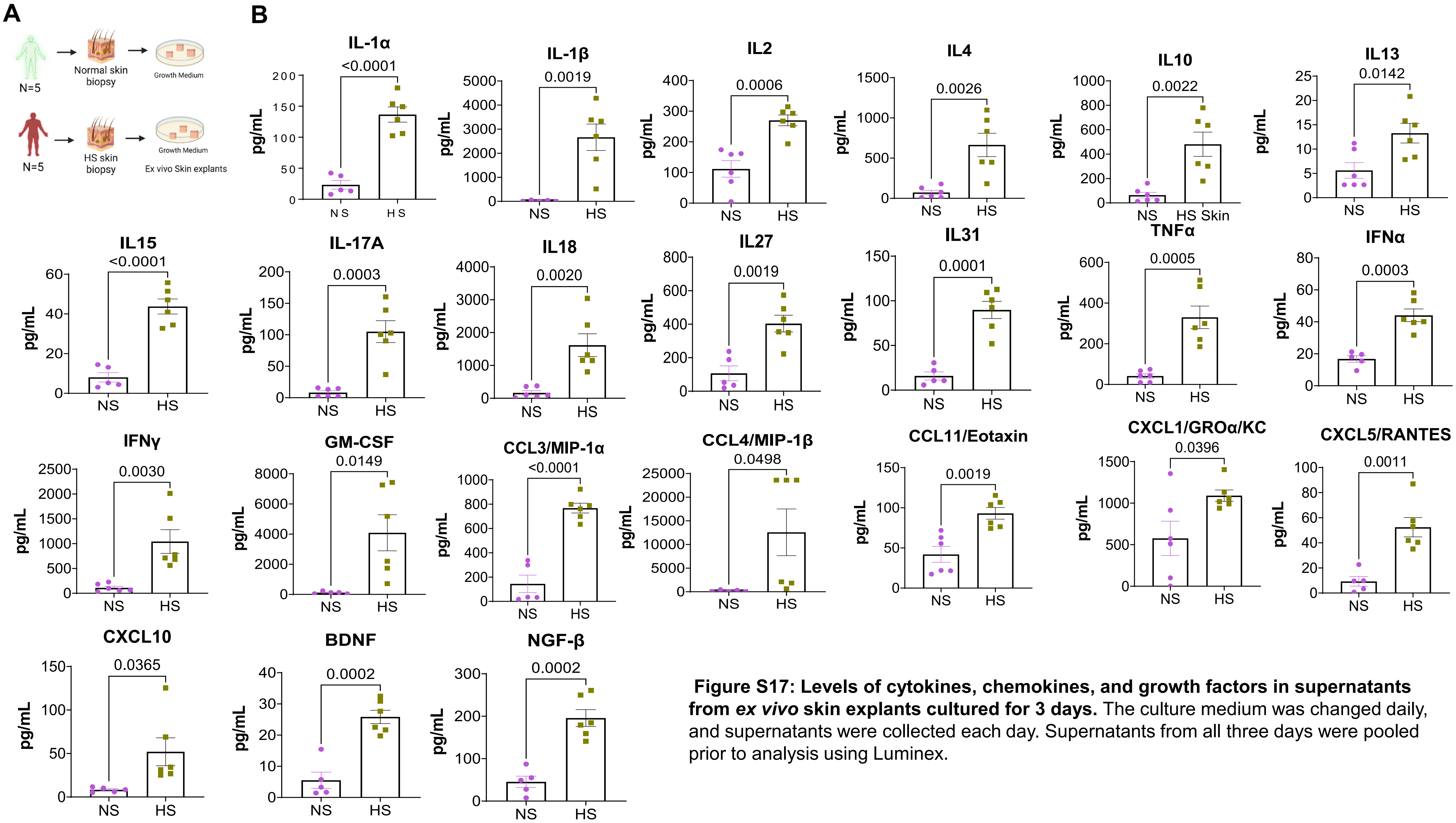

**Figure S17: Levels of cytokines, chemokines, and growth factors in supernatants from *ex vivo* skin explants cultured for 3 days.** The culture medium was changed daily, and supernatants were collected each day. Supernatants from all three days were pooled prior to analysis using Luminex.

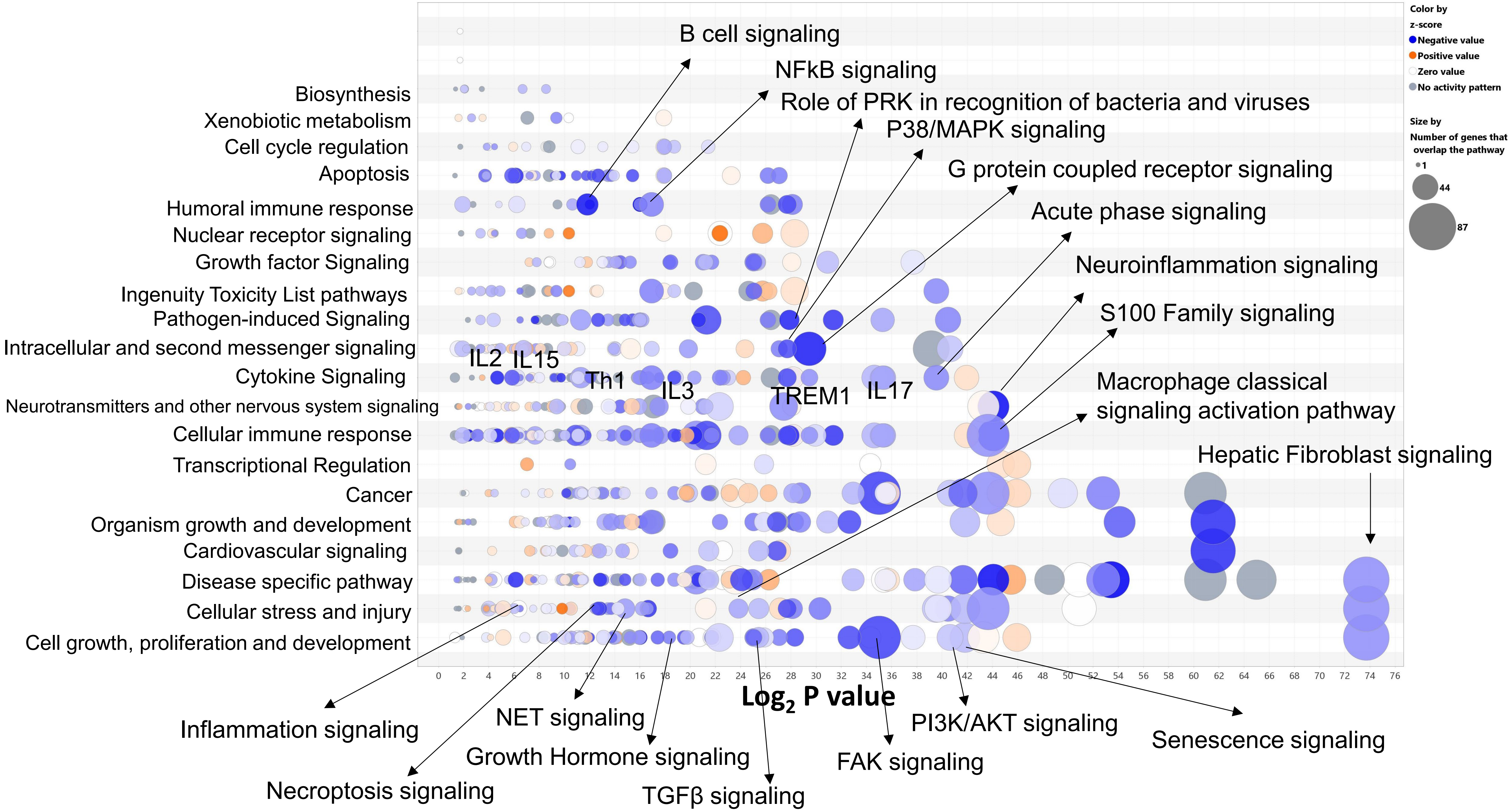

**Figure S18:** Bubble chart from IPA analysis of bulk qPCR data from rapamycin-treated HS skin explants.

### Autophagy Network

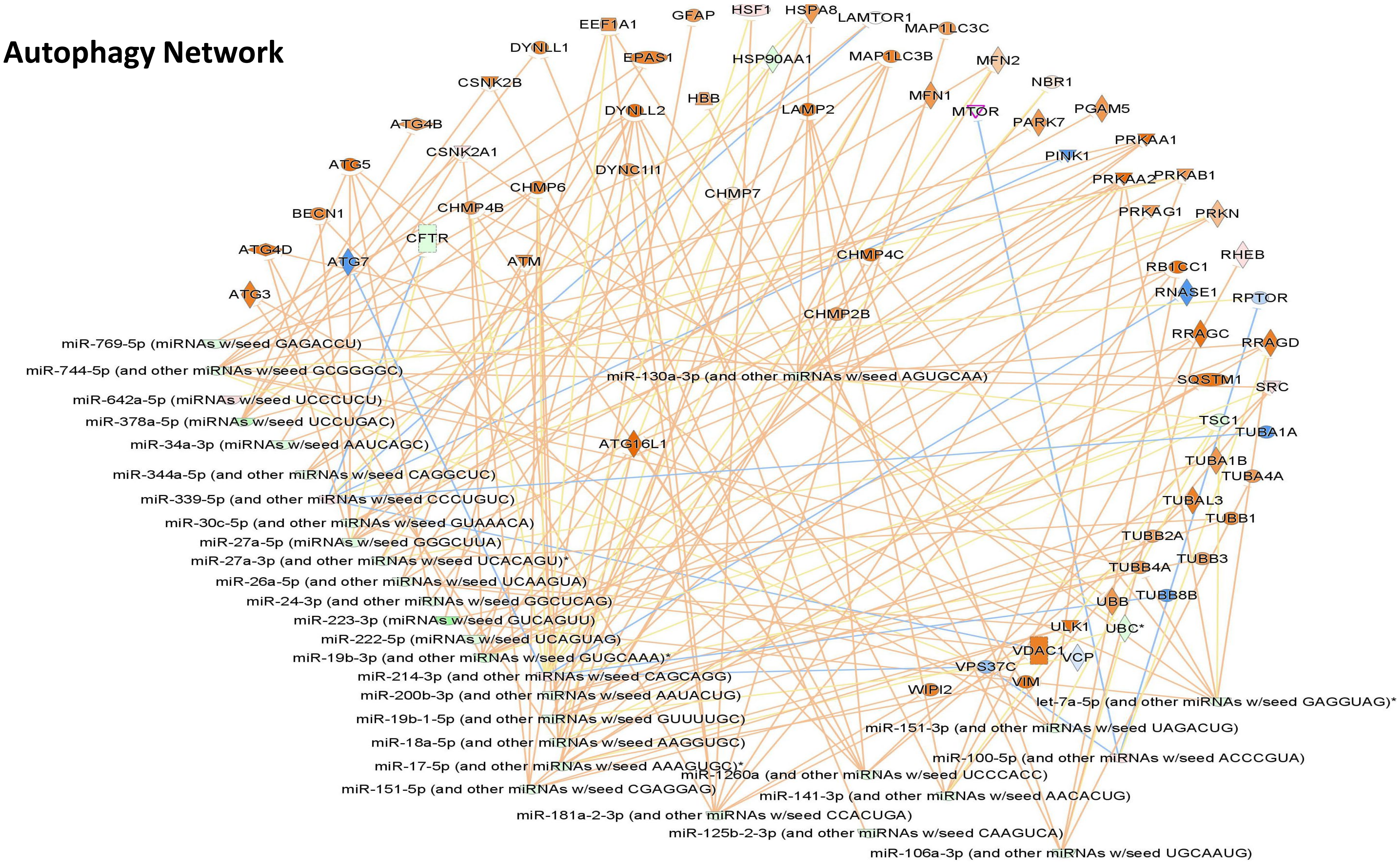

**Figure S19:** IPA analysis of the miRNA–mRNA interactome showing rapamycin-induced autophagy network. miRNA–mRNA autophagy regulatory network was constructed by integrating OpenArray TaqMan-based miRNA profiling with mRNA expression data from Inflammation and Signal Transduction panels.

Table S1: Demographic characteristics and tissue sources of HS and control skin samples, with a summary of samples used across the different experimental analyses.

| | Serial Number | Race | Sex | Tissue Location | Experiments |
|---|---|---|---|---|---|
| **1** | HS # 1 | Black | F | R-Breast | M & L, PRO, KINO, QPCR, WB |
| **2** | HS # 2 | Black | F | Armpit | M & L, QPCR, WB |
| **3** | HS # 3 | Black | M | gluteal | M & L, PROT, KINO, QCR, IF, WB |
| **4** | HS # 4 | Black | F | groin | M & L, PROT, KINO, QPCR, WB |
| **5** | HS # 5 | Black | M | groin | M & L, PROT, KINO, QPCR, IF, WB |
| **6** | HS # 6 | Black | M | Back | M & L, QPCR, WB |
| **7** | HS # 7 | Black | M | Armpit | M & L, PROT, KINO, QPCR, IF, WB |
| **8** | HS # 8 | Black | F | Abdominal | M & L, QPCR, WB |
| **9** | HS # 9 | Black | M | Groin | M & L, WB |
| **10** | HS # 10 | Black | M | Axilla | M & L |
| **11** | HS # 11 | Black | M | Axilla | M & L, EXCL-ND, LUM |
| **12** | HS # 12 | Black | M | Buttock | M & L, EXCL-ND, LUM |
| **13** | HS # 13 | Black | F | Arm | M & L, EXCL-ND, LUM |
| **14** | HS # 14 | Black | F | Armpit | M & L, EXCL-ND, LUM |
| **15** | HS # 15 | Black | F | Axilla | EXCL-ND, IF, LUM |
| **16** | HS # 16 | Black | F | Unknown | EXCL, DRUG, ARRAY, LUM |
| **17** | HS # 17 | Black | F | Unknown | EXCL, DRUG, ARRAY, LUM |
| **18** | HS #18 | Black | F | Unknown | EXCL, DRUG, ARRAY, LUM |
| **19** | HS # 19 | Black | F | Axillar | EXCL, DRUG, LUM |
| **20** | HS #20 | Black | M | Groin | EXCL, DRUG, LUM, IF |
| **1** | NS # 1 | Caucasian | F | Breast | M & L, QPCR, IF, WB |
| **2** | NS # 2 | Black | F | Armpit | M & L, PROT, KIN, QPCR, WB |
| **3** | NS # 3 | Black | M | Back | M & L, PROT, KIN, QPCR, IF, WB |
| **4** | NS # 4 | Black | F | Breast | M & L, QPCR |
| **5** | NS # 5 | Black | M | Abdominal | M & L, PROT, KIN, QPCR, IF, WB |
| **6** | NS # 6 | Black | F | Breast | M & L PROT, KIN, QPCR, WB |
| **7** | NS # 7 | Black | F | Breast | M & L, QPCR |
| **8** | NS # 8 | Black | F | Breast | M & L, QPCR, EXCL-ND, LUM |
| **9** | NS # 9 | Black | M | Breast | M & L, EXCL-ND, LUM |
| **10** | NS # 10 | Hispanic | M | Abdominal | M & L, EXCL-ND, IF, LUM |
| **11** | NS # 11 | Black | F | Brest | EXCL-ND, LUM |
| **12** | NS # 12 | Black | F | Breast | EXCL-ND, LUM |
| **13** | NS # 13 | Black | F | Breast | EXCL, DRUG, ARRAY, LUM |
| **14** | NS # 14 | Black | F | Breast | EXCL, DRUG, ARRAY, LUM |

| | | | | | |
|---|---|---|---|---|---|
| **15** | NS # 15 | Caucasian | F | Breast | EXCL, DRUG, ARRAY, LUM |
| **16** | NS # 16 | Caucasian | F | Abdominoplasty | EXCL, DRUG, LUM |
| **17** | NS # 17 | Caucasian | F | Breast | EXCL, DRUG, LUM |

HS = Lesional skin from Hidradenitis suppurative patients
NS= Normal Skin
M & L= Metabolomics and Lipidomics
PRO= Proteomics and phospho-proteomics
KIN= Kinomics
QPCR, TaqMan based qRT-PCR
EXCL= *ex vivo skin explant culture*
*EXCL-ND= ex vivo skin explant culture (without any drug-treatment)*
*ARRAY= TaqMan based Openarray*
*LUM= Luminex Multiplexing*
*IF= Immunofluorescences Confocal microscopy*

Note: All HS patients had late-stage Hurley II/III disease.

**Table S2:** List of antibodies used for each IHC or immunofluorescence microscopy and TaqMan primers used for qPCR.

| Antibodies | Cat No | Company | Dilution | Assay |
|---|---|---|---|---|
| NDP52 | NBP2-19499 | NOVUS Biologicals | 1:1000 | WB |
| ATG16L1 | 8089T | Cell Signaling Technology | 1:1000 | WB |
| ATG12 | 4180S | Cell Signaling Technology | 1:1000 | WB |
| ATG5 | 12994S | Cell Signaling Technology | 1:1000 | WB |
| ATG3 | 3415S | Cell Signaling Technology | 1:1000 | WB |
| Beclin-1 | 3495S, | Cell Signaling Technology | 1:1000 | WB |
| P62 | ab109012 | Abcam | 1:1000 | WB |
| LC3A/B | 4108S | Cell Signaling Technology | 1:1000 | WB |
| pERK | 4376S | Cell Signaling Technology | 1:1000 | WB |
| ERK | ab17942 | Cell Signaling Technology | 1:1000 | WB |
| pNFkB | ab30623 | Abcam | 1:500 | WB |
| NRF2 | ab76026 | Abcam | 1:500 | WB |
| HO1 | 5853S | Cell Signaling Technology | 1:500 | WB |
| 4-Hydroxynonenal | ab46545 | Abcam | 1:500 | WB |
| TRPV1 | ab6166 | Abcam | 1:500 | WB |
| TRPV2 | ab6183 | Abcam | 1:500 | WB |
| TRPV4 | ab14455 | Abcam | 1:500 | WB |
| mTOR | 2983S | Cell Signaling Technology | 1:500 | WB |
| Phospho-AMPKα | ab23875 | Abcam | 1:500 | WB |

| | | | | |
|---|---|---|---|---|
| pMTOR (ser 2448) | 5536 | Cell Signaling Technology | 1:500 | WB |
| Anti-β-Actin (ACTB) Antibody | A5441 | Millipore Sigma | 1:3000 | WB |
| P62 | ab56416 | Abcam | 1:100 | IF |
| CD68 | 916106 | Bioligand | 1:100 | IF |
| NLRP3 | 27458-1 | Proteintech | 1:100 | IF |
| NLRP3 | MAB7578 | R&D Systems | 1:100 | IF |
| NLRP1 | ab36852 | Abcam | 1:150 | IF |
| S100A8/A9 | ab22506 | Abcam | 1:100 | IF |
| Anti-CD2 antibody | AF1856 | R&D Systems | 1:250 | IF |
| Anti-NCAM-1/CD56 Antibody | 13-0567-82 | eBioscience | 1:50 | IF |
| Anti-mTOR (phospho S2448) antibody [EPR426(2)] | ab131538 | Abcam | 1:100 | IF |
| ZKSCAN3 | ab187866 | Abcam | 1:100 | IF |
| CK14 | MA5-11599 | Thermofischer | 1:200 | IF |

**TaqMan Assay:**

| S. No. | Gene Name | Assay ID | Company |
|---|---|---|---|
| 1 | NLRP3 | Hs00918082_m1 | Thermofisher Scientific |
| 2 | PYCARD | Hs00203118_m1 | Thermofisher Scientific |
| 3 | CASP1 | Hs00354836_m1 | Thermofisher Scientific |
| 4 | ACTB | Hs99999903_m1 | Thermofisher Scientific |
